# Spatial control of secretory vesicle targeting by the Ync13–Rga7– Rng10 complex during cytokinesis

**DOI:** 10.1101/2025.05.13.653810

**Authors:** Sha Zhang, Davinder Singh, Yi-Hua Zhu, Katherine J. Zhang, Alejandro Melero, Sophie G. Martin, Jian-Qiu Wu

## Abstract

Cytokinesis requires precise coordination of contractile-ring constriction, vesicle trafficking and fusion to the plasma membrane, and extracellular matrix assembly/remodeling at the cleavage furrow to ensure faithful cell division and maintain cell integrity. These processes and proteins involved are broadly conserved across eukaryotes, yet molecular mechanisms controlling the spatiotemporal pathways of membrane trafficking remain poorly understood. Here, using fission yeast genetics, microscopy, and in vitro binding assays, we identify a conserved module including the Munc13 protein Ync13, F-BAR protein Rga7, and coiled-coil protein Rng10 to be critical for precise and selective vesicle targeting during cytokinesis. The module specifically recruit the TRAPP-II but not exocyst complex to tether vesicles containing the glucan synthases Bgs4 and Ags1 along the cleavage furrow. Ync13 subsequently interacts with the SM protein Sec1 for vesicle fusion. Mutations in this pathway disrupt septum integrity and lead to cell lysis. Our work provides key insights into how membrane trafficking is tightly controlled to maintain cell integrity during cytokinesis.

## Introduction

Cytokinesis is a highly conserved cellular process occurring at the late stage of the cell division cycle, resulting in the generation of two daughter cells. From fungi to humans, this process involves coordinated actions of division site selection, actomyosin contractile ring assembly and constriction, plasma membrane deposition and invagination at the cleavage furrow, and extracellular matrix formation or remodeling (Bement, 2002; Feierbach and Chang, 2001; Gerien and Wu, 2018; Robinson and Spudich, 2000; Wang et al., 2016; Wu et al., 2003). In fungal cells including the fission yeast *Schizosaccharomyces pombe*, two layers of new plasma membranes and a division septum form behind the constricting ring during cytokinesis. The septum is a three-layered cell wall structure that consists of a middle primary septum flanked by secondary septa on each side.

The septum is primarily constructed by three essential transmembrane glucan synthases. The β-Glucan synthase Bgs1/Cps1 synthesizes the linear β(1,3)-glucan for the primary septum (Cortes et al., 2002; Cortes et al., 2007; Liu et al., 1999), while the β-glucan synthase Bgs4/Cwg1 and the α-glucan synthase Ags1/Mok1 are mainly involved in the formation of the secondary septum (Calonge et al., 2000; Cortes et al., 2005; Cortes et al., 2012; Katayama et al., 1999; Longo et al., 2022). Once the septum is complete and mature, daughter cells separate via digesting the primary septum mainly by the glucanases Eng1 and Agn1 (Baladrón et al., 2002; Dekker et al., 2004; Martin-Cuadrado et al., 2003). Proper plasma membrane deposition, septum formation, and cell separation are essential for maintaining cell integrity and viability during cytokinesis, especially due to the high cellular turgor pressure in fungal cells (Barba et al., 2008; Cansado et al., 2021; Roncero et al., 2021; Roncero and Sanchez, 2010). The F-BAR protein Cdc15, transmembrane protein Sbg1, the Transport Particle Protein II (TRAPP-II) complex, Rho1 GTPase, and other proteins help recruit and/or activate Bgs1 behind the contractile ring to build the primary septum (Arasada and Pollard, 2014; Arellano et al., 1997; Bhattacharjee et al., 2023; Cortes et al., 2002; Davidson et al., 2016; Garcia et al., 2006a; Nakano et al., 1997; Ren et al., 2015; Roberts-Galbraith et al., 2010; Sethi et al., 2016; Wang et al., 2016). Cdc15 binds to the plasma membrane through its F-BAR domain and helps deliver Bgs1 to the plasma membrane while paxillin Pxl1 mediates the interaction of Bgs1 to the contractile ring (Arasada and Pollard, 2014; Cortés et al., 2015; Roberts-Galbraith et al., 2009). Septins, the anillin Mid2, the exocyst complex, the Rho3 and Rho4 GTPases, and Rho4 GEF Gef3 concentrate the glucanase Eng1 to the rim of the division plane for daughter-cell separation (Alonso-Nunez et al., 2005; Berlin et al., 2003; Martin-Cuadrado et al., 2005; Pérez et al., 2015; Santos et al., 2005; Singh et al., 2024; Tasto et al., 2003; Wang et al., 2003; Wang et al., 2015). In addition, the TRAPP-II complex is also important for Eng1’s localization at the centroid of the division plane (Wang et al., 2016). However, the mechanisms that control the precise spatiotemporal localizations of Bgs4 and Ags1 on the plasma membrane at the division plane for secondary septum formation remain poorly understood.

The coordination of exocytosis and endocytosis is essential for successful cytokinesis. Exocytosis delivers the proteins and membranes needed for furrow ingression and extracellular matrix (including cell wall) formation or remodeling to the division site (Baladrón et al., 2002; Dekker et al., 2004; Martin-Cuadrado et al., 2003; Martin-Cuadrado et al., 2005; Wang et al., 2002). Exocytosis, a highly regulated process, involves several sequential stages: vesicle delivery/trafficking, tethering and docking onto the plasma membrane, priming of the fusion machinery, and membrane fusion via SNARE complex assembly (Gerien and Wu, 2018; Grote et al., 2000; Hsu et al., 2004; Liu and Guo, 2012; Luo et al., 2014; Weber-Boyvat et al., 2013). In fission yeast cytokinesis, the TRAPP-II complex recognizes and tethers secretory vesicles along the whole cleavage furrow, while the octomeric tethering complex exocyst mainly tethers vesicles that contain glucanases Eng1 and Agn1 and other cargos at the rim of the division plane although it also functions along the furrow (Singh et al., 2024; Wang et al., 2002; Wang et al., 2003; Wang et al., 2016). During mammalian synaptic exocytosis, after vesicle tethering, three SNARE proteins—synaptobrevin-2 (Syb2) on synaptic vesicles, and syntaxin-1 (Syx1) and SNAP-25 (SN25) on the plasma membrane form a ternary trans-SNARE complex (Jahn and Scheller, 2006; Sutton et al., 1998; Weber et al., 1998). This complex brings the vesicle and plasma membrane into close proximity to facilitate membrane fusion. Vesicle tethering and fusion are regulated by Rab GTPases and several other proteins including: the Sec1/Munc18 (SM) family protein Munc18-1, which initially locks Syx1 in a closed conformation to inhibit SNARE complex assembly; and the Munc13/UNC-13 family protein Munc13-1, which catalyzes the transition from the Munc18-1/Syx1 complex to the SNARE complex in the presence of SN25 and Syb2 (Bhaskar et al., 2024; Gerien et al., 2020; Guan et al., 2008; Leitz et al., 2024; Shu et al., 2020). In *S. pombe*, Ync13 is the homolog of Munc13 and UNC-13. Ync13 localizes to cell tips during interphase and the plasma membrane at the cleavage furrow during cytokinesis (Zhu et al., 2018). Deletion of Ync13 results in defective exocytosis, impaired endocytosis, uneven distribution of cell wall enzymes at the division site, and extensive cell lysis during cell separation (Zhu et al., 2018). However, the regulatory mechanisms of Ync13 in membrane dynamics, its binding partners, and the exact cause of cell lysis upon its deletion remain unclear.

Endocytosis also plays essential and dynamic roles in cytokinesis, particularly in membrane remodeling, signal regulation, and recycling membrane proteins. Cells internalize membrane lipids and proteins from the plasma membrane mainly through the clathrin-mediated endocytosis from growth and division sites (Feng et al., 2002; Gachet and Hyams, 2005; Kaksonen et al., 2005; Kukulski et al., 2012a; Lemière et al., 2021; Pearse, 1976; Weinberg and Drubin, 2012; Yang et al., 1997). During cytokinesis, endocytosis helps clear old/inactive or excess membrane proteins and recycle materials to be reused for septum formation or membrane expansion (Feng et al., 2002; Gachet and Hyams, 2005; Gerien and Wu, 2018). Thus, endocytosis works hand-in-hand with exocytosis to maintain the proper levels of glucan synthases and other cell wall modifying enzymes needed for septum formation. Yeast endocytosis is actin dependent and many endocytic mutants in genes such as the endocytic adaptor *ede1*, clathrin light chain *clc1*, fimbrin *fim1*, and *cdc15* result in furrow ingression defects, cell wall abnormalities, and cytokinesis failure (Arasada and Pollard, 2011; de León et al., 2013; Gachet and Hyams, 2005; Laporte et al., 2012; Martin-Garcia et al., 2014; Skau and Kovar, 2010; Suzuki et al., 2012; Wu et al., 2001).

Rga7 is a Rho2 GAP in fission yeast that plays crucial roles in cytokinesis (Arasada and Pollard, 2015; Liu et al., 2016; Liu et al., 2019; Martin-Garcia et al., 2014). It contains an F-BAR domain necessary for its membrane association and a Rho GAP domain at its C-terminus. Rga7 localizes to the plasma membrane at the cell tips during interphase and relocates to the division site during cytokinesis, a process dependent on its interaction with the coiled-coil protein Rng10 and membrane lipids (Arasada and Pollard, 2015; Liu et al., 2016; Liu et al., 2019; Martin-Garcia et al., 2014). Without Rng10, Rga7 essentially disappears from the division site, leading to defective septum formation and cell lysis (Liu et al., 2016; Liu et al., 2019). Rga7 collaborates with F-BAR proteins Cdc15, Imp2, Fic1, and Pxl1 to maintain actomyosin ring stability and ensure successful septum formation and separation (Demeter and Sazer, 1998; Ge and Balasubramanian, 2008; Martin-Garcia et al., 2014; Pinar et al., 2008; Roberts-Galbraith et al., 2009). Similar to Ync13, Rga7 and Rng10 collaboratively regulate the accumulation and dynamics of glucan synthases (Liu et al., 2016). Rga7 is reported to facilitate the trafficking of β-glucan synthase Bgs4 from the Golgi to the plasma membrane (Arasada and Pollard, 2015). Cells lacking Rga7 show defects similar to those observed in *bgs4* and *ync13* mutant cells (Liu et al., 2016; Munoz et al., 2013; Zhu et al., 2018). However, the relationships between Ync13, Rga7/Rng10, the TRAPP-II complex, and Sec1 in cytokinesis were unknown.

In this study, we aimed to map out some key spatiotemporal pathways for plasma membrane deposition and septum formation during cytokinesis using fission yeast as a model system by mistargeting proteins to mitochondria, co-immunoprecipitation, in vitro binding assays, genetic and cellular methods, electron microscopy, and live-cell confocal microscopy. We find that Ync13 regulates exocytosis by interacting with Rga7-Rng10, the SM protein Sec1, and the TRAPP-II complex. The Ync13-Rga7-Rng10 module selectively controls the accumulation and distribution of glucan synthases involved in secondary septum formation, such as Bgs4 and Ags1; rather than Bgs1, which is essential for the primary septum assembly.

Consistently, the secondary septum is defective in *ync13* mutants, which leads to cell lysis during daughter-cell separation. Collectively, we find that the Ync13-Rga7-Rng10 module and the TRAPP-II complex are central players for plasma membrane dynamics and secondary septum formation during cytokinesis.

## Results

### Mapping the physical interactions among the key cytokinetic proteins involved in plasma membrane deposition and septum formation by ectopic mistargeting

Unlike the proteins in the contractile ring and its precursor nodes (Laplante et al., 2016; Laporte et al., 2011; McDonald et al., 2017; Padmanabhan et al., 2011; Wu et al., 2006), the physical interactions among the proteins involved in plasma membrane deposition/expansion and septum formation are poorly understood. These proteins are essential for plasma membrane and septum integrity, exocytosis, and endocytosis. In this study, we used the fission yeast *S. pombe* as a model system to test the physical interactions among the key proteins by first mistargeting mEGFP, GFP, or mECitrine tagged proteins to mitochondria using outer mitochondrial membrane protein Tom20 (Yamamoto et al., 2011) tagged with GFP-binding protein (GBP) nanobody, then examined if tdTomato, mCherry, or RFP tagged proteins were also mistargeted to mitochondria. All the tagged genes are the sole copies of the genes in the cells. Our and others’ previous studies have shown this strategy is highly efficient at detecting protein physical interactions (Liu et al., 2016; Liu et al., 2019; Longo et al., 2022; Luo et al., 2014; Wang et al., 2016; Yamamoto et al., 2011). As shown previously (Liu et al., 2016; Rothbauer et al., 2008; Schornack et al., 2009), GBP does not bind to tdTomato, mCherry, or RFP; and no signal bleed through between the green and red channels were detected (Fig. S1).

We started with Ync13, Rga7, and Rng10 because they perfectly colocalized on the plasma membrane at cell tips and the division site (Fig. 1 A), and their mutations lead to cell lysis during daughter-cell separation (Arasada and Pollard, 2015; Liu et al., 2016; Liu et al., 2019; Martin-Garcia et al., 2014; Zhu et al., 2018). We first mistargeted Rga7 or Rng10 to the mitochondria using Tom20-GBP. The ectopically targeted Rga7 and Rng10 were both able to recruit Ync13-tdTomato to the mitochondria (Fig. 1 B). We detected mitochondrial localization of Ync13 in all Tom20-GBP Rga7-mEGFP (n = 120 cells) and Tom20-GBP Rng10-mEGFP expressing cells (n = 200 cells), except in some unhealthy cells that Ync13 diffused in the whole cytoplasm, which could be due to the side effects (besides cell lysis which is obvious in the DIC channel) of mislocalization of Rga7, Rng10, and Ync13 (Fig. 1 B). Mistargated proteins concentrated into clusters instead of over the whole mitochondria (Fig. 1 B). In addition, the ectopically targeted Ync13 recruited the Rga7 and Rng10 to the mitochondria (Fig. S2 A). In this assay, Ync13 was overexpressed using the *3nmt1* promoter under repressed condition (in medium with thiamine) because Ync13’s native level was too low to spread sufficiently to the abundant Tom20-GBP.

**Figure 1.**
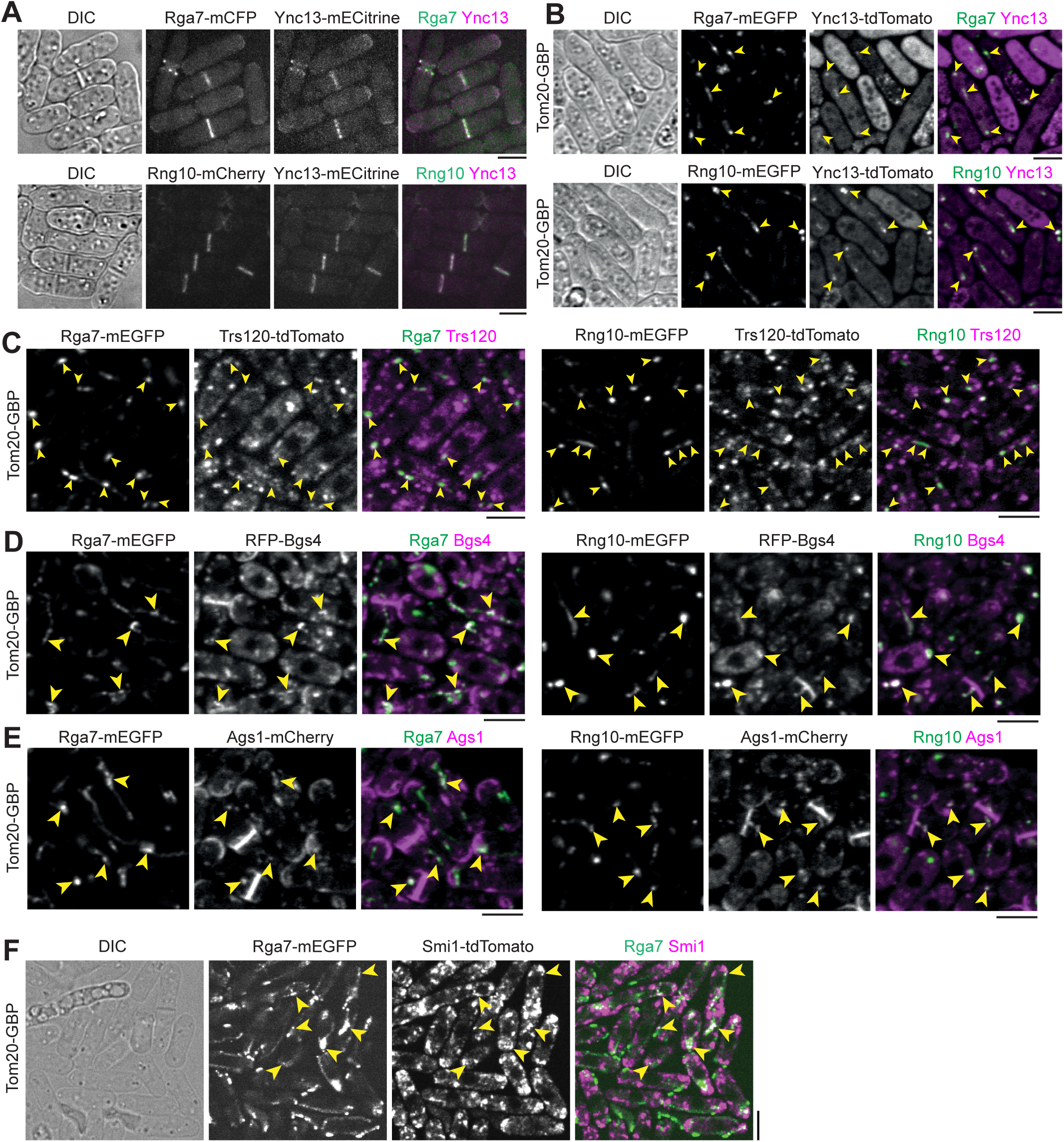
Physical interactions among the key cytokinetic proteins in plasma membrane deposition and septum formation revealed by ectopic mistargeting to mitochondria by Tom20-GBP. Arrowheads mark examples of colocalization at mitochondria. **(A)** Ync13 colocalizes with Rga7 and Rng10 at cell tips and the division site. **(B-F)** Tom20-GBP can ectopically mistarget pairs of proteins with Rga7/Rng10-mEGFP and tdTomato/RFP/mCherry tagged proteins to mitochondria. **(B)** Rga7/Rng10-Ync13. **(C)** Rga7/Rng10-Trs120. **(D)** Rga7/Rng10-Bgs4. **(E)** Rga7/Rng10-Ags1. **(F)** Rga7-Smi1. Bars, 5 μm.

**Table 1.** Summary of Tom20-GBP mistargeting assays.

| Protein 1 | Protein 2 | Protein 2<br>mislocalized to<br>mitochondria (Y/N) | Figure |
| --- | --- | --- | --- |
| Rga7-mEGFP | Ync13-tdTomato | Y | Fig 1 B |
| Rng10-mEGFP | Ync13-tdTomato | Y | Fig 1 B |
| Rga7-mEGFP | Trs120-tdTomato | Y | Fig 1 C |
| Rng10-mEGFP | Trs120-tdTomato | Y | Fig 1 C |
| Rga7-EGFP | RFP-Bgs4 | Y | Fig 1 D |
| Rng10-mEGFP | RFP-Bgs4 | Y | Fig 1 D |
| Rga7-mEGFP | Ags1-mCherry | Y | Fig 1 E |
| Rng10-mEGFP | Ags1-mCherry | Y | Fig 1 E |
| Rga7-mEGFP | Smi1-tdTomato | Y | Fig 1 F |
| 3nmt1-mECitrine-Ync13 | Rga7-mCherry | Y | Fig S2 A |
| 3nmt1-mECitrine-Ync13 | Rng10-mCherry | Y | Fig S2 A |
| 3nmt1-mECitrine-Ync13 | Sec1-tdTomato | Y | Fig S2 B |
| 3nmt1-mECitrine-Ync13 | RFP-Bgs4 | Y | Fig S2 H |
| 3nmt1-mECitrine-Ync13 | Ags1-mCherry | Y | Fig S2 H |
| Rga7-FBD-mEGFP | Ync13-tdTomato | Y | Fig S3 B |
| Rga7(ΔF-BAR)-mEGFP | Ync13-tdTomato | Y | Fig S3 C |
| mECitrine-Rng10-(201-1038) | Ync13-tdTomato | Y | Fig S3 E |
| Rng10(1-750)-mEGFP | Ync13-tdTomato | Y | Fig S3 F |
| mECitrine-Rng10(751-1038) | Ync13-tdTomato | Y | Fig S3 G |
| 3nmt1-mECitrine-Ync13 | Sec3-mCherry | N | Fig S2 C |
| 3nmt1-mECitrine-Ync13 | Ede1-mCherry | N | Fig S2 D |
| 3nmt1-mECitrine-Ync13 | Fim1-mCherry | N | Fig S2 E |
| 3nmt1-mECitrine-Ync13 | Clc1-mCherry | N | Fig S2 F |
| Rga7-mEGFP | Sec3-tdTomato | N | Fig S2 G |
| Rng10-mEGFP | Sec3-tdTomato | N | Fig S2 G |
| Rga7-mEGFP | tdTomato-Bgs1 | N | Fig S2 I |
| Rng10-mEGFP | tdTomato-Bgs1 | N | Fig S2 I |
| Rng10-(1-200)-mEGFP | Ync13-tdTomato | N | Fig S3 D |

Because *ync13* mutants are defective in both exocytosis and endocytosis (Zhu et al., 2018), we tested if Ync13 could mistarget key proteins in exocytic and endocytic pathways to mitochondria by Tom20-GBP. The ectopically targeted mECitrine-Ync13 was able to recruit the Sec1/Munc18 (SM) family protein Sec1-tdTomato to the mitochondria (Fig. S2 B), suggesting Ync13 may function as a priming factor for SNARE complex assembly and/or vesicle tether during exocytosis similar to its animal homolog Munc13/UNC-13 (Bhaskar et al., 2024; Dulubova et al., 2005; Guan et al., 2008; Khodthong et al., 2011; Li et al., 2011; Ma et al., 2011; Pei et al., 2009; Shu et al., 2020). However, Ync13 could not recruit the following proteins to mitochondria: the exocyst subunit Sec3 (Fig. S2 C); or endocytic proteins early coat marker Eps15 protein Ede1 (Suzuki et al., 2012), actin crosslinker fimbrin Fim1 (Nakano et al., 2001; Wu et al., 2001), and clathrin light chain Clc1 (de León et al., 2013) (Fig. S2, D-F). Thus, Ync13 interacts with the SNARE-binding protein Sec1 that is involved in exocytosis, but not with the exocyst or various proteins in endocytosis.

We next tested if Rga7 or Rng10 can mistarget TRAPP-II vesicle tether and secretory cargos to mitochondria by Tom20-GBP. Both Rga7-mEGFP and Rng10-mEGFP recruited Trs120-tdTomato (Cai et al., 2005; Choi et al., 2011), but not the exocyst subunit Sec3 (He and Guo, 2009), to mitochondria (Fig. 1 C and Fig. S2 G). These data suggest that Rga7 and Rng10 selectively interact with the TRAPP-II complex to promote vesicle tethering during exocytosis along the cleavage furrow, rather than the exocyst complex, which is more concentrated at the rim of the division plane (Singh et al., 2024; Wang et al., 2016).

Mistargeted Rga7-mEGFP and Rng10-mEGFP ectopically recruited RFP-Bgs4 to mitochondria in ∼100% of Tom20-GBP Rga7-mEGFP or Tom20-GBP Rng10-mEGFP cells (Fig. 1 D, n > 100 cells), and Ags1-mCherry to mitochondria in ∼90% of Tom20-GBP Rga7-mEGFP or Tom20-GBP Rng10-mEGFP cells (Fig. 1 E, n > 100 cells). Consistently, Rga7 mistargeted tdTomato tagged Smi1, which is an adaptor for Bgs4 (Longo et al., 2022), to mitochondria in cells expressing Tom20-GBP Rga7-mEGFP (Fig. 1 F). Similarly, Ync13-mECitrine could mistarget Bgs4 and Ags1 to mitochondria (Fig. S2 H). Surprisingly, neither ectopically targeted Rga7 nor Rng10 recruited Bgs1 to mitochondria (Fig. S2 I), suggesting the interactions of Rga7 and Rng10 with the glucan synthases are selective.

Ectopic mistargeting of Ync13-tdTomato to mitochondria using Rga7 or Rng10 truncations further supported that Rga7, Rng10, and Ync13 interact with each other (Fig. S3). Except Rng10-(1-200) (Fig. S3 D), all other Rga7 and Rng10 truncations (Fig. S3) mistargeted Ync13-tdTomato to mitochondria, although less efficiently than the full-length proteins. Based on what we know about Rga7 and Rng10 interactions and localization interdependency (Liu et al., 2016; Liu et al., 2019), the data suggest that Rga7, Rng10, and Ync13 have multivalent interactions with each other. Collectively, these mistargeting data suggest Rng10, Rga7, and Ync13 form a protein complex, which recruits the TRAPP-II complex and Sec1 but not the exocyst to the plasma membrane for vesicle tethering, SNARE complex assembly, and fusion. The main proteins recruited by the Rng10/Rga7/Ync13 complex are secretory vesicles containing cargos such as the glucan synthases Bgs4 (and its adaptor Smi1) and Ags1, but not Bgs1 or the endocytic machinery.

### Rga7 physically interacts with Ync13, Bgs4, and Smi1

We used co-immunoprecipitation (Co-IP) and in vitro binding assays to confirm some of the major interactions revealed by the mistargeting experiments. It is known that Rng10 directly interacts with Rga7 and both proteins interact with membrane lipids (Liu et al., 2016; Liu et al., 2019). The F-BAR domain of Rga7 interacts with Rng10 C-terminal amino acids 751-1038 (Liu et al., 2019). Smi1 interacts with Bgs4 and is important for Bgs4 localization (Longo et al., 2022). Most other interactions suggested by the mistargeting assays had not been tested. For Co-IP assays, all the proteins were expressed under their native promotors. Consistent with the mistargeting results (Figs. 1, S2 and S3), Rga7-13Myc was coimmunoprecipitated from fission yeast cell extracts by Ync13-mECitrine (Fig. 2 A), GFP-Bgs4 (Fig. 2 B), and Smi1-mEGFP (Fig. 2 C), indicating these proteins do interact or form big protein complexes. Moreover, Smi1-13Myc also coimmunoprecipitated by Rga7-mEGFP (Fig. 2 C). Furthermore, when Ync13-3FLAG expressed at its native chromosomal locus and under endogenous promotor was immunoprecipitated from *S. pombe* cell extracts, we identified Rga7 and Rng10 reproducibly among Ync13’s binding partners by mass spectrometry analyses. These results suggest that Rga7, Rng10, and Ync13 form a protein complex.

**Figure. 2.**
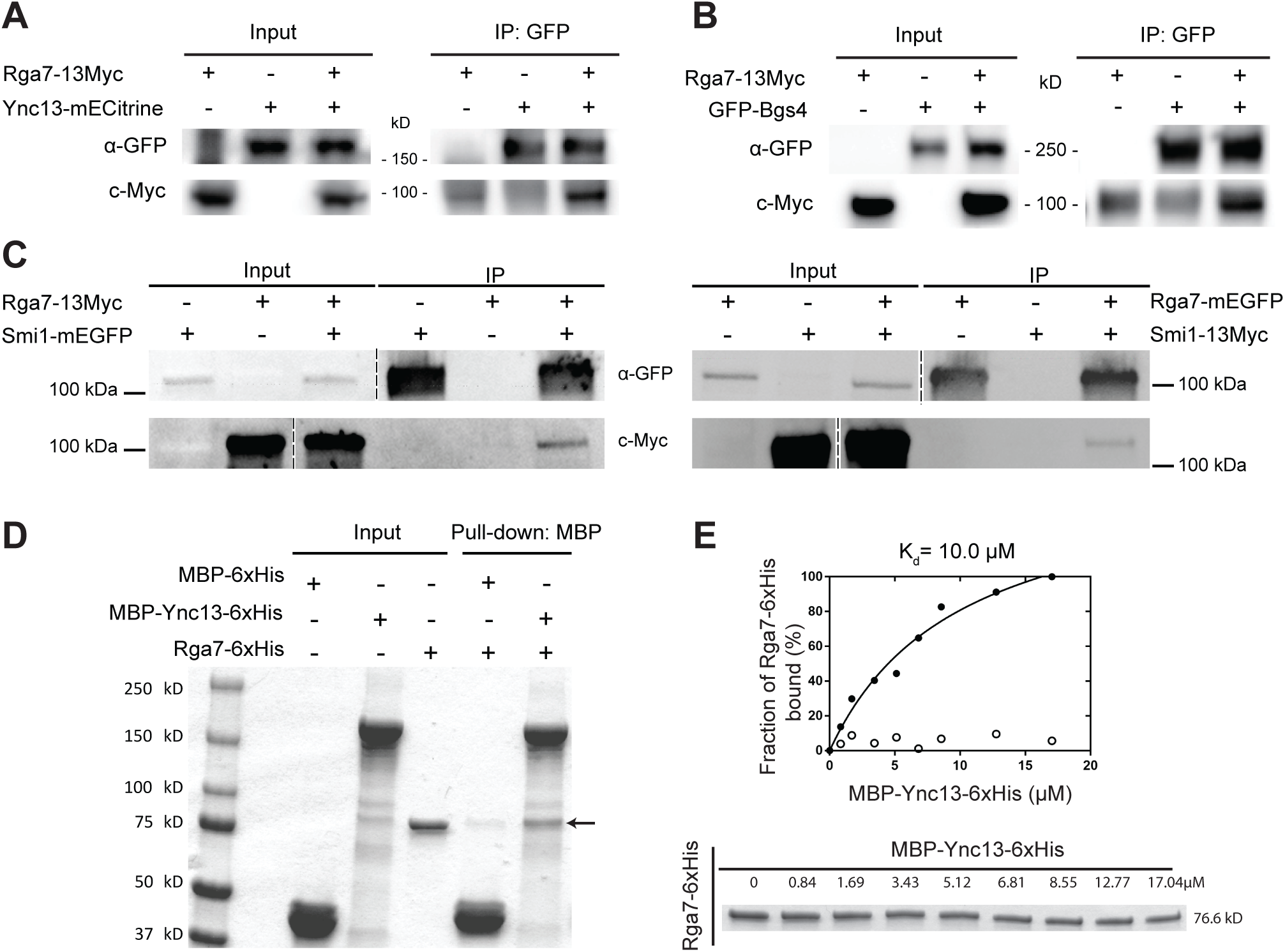
Confirmation of Rga7’s physical interactions with Ync13, Bgs4, Smi1 by co-IP and/or in vitro binding assays. **(A-C)** Rga7-13Myc coimmunoprecipitates with Ync13 **(A)**, Bgs4 **(B)**, and Smi1 **(C, left)** from *S. pombe* cell extracts using antibodies against GFP. **(C, right)** Smi1-13Myc coimmunoprecipitates with Rga7. The vertical dashed lines mark the positions of protein ladders that were excised out. **(D)** In vitro binding of Rga7 and Ync13 using purified proteins. Bead bound MBP-Ync13-6xHis or MBP-6xHis control was incubated with Rga7-6xHis. The arrow marks the Rga7-6xHis band. **(E)** Supernatant depletion assay to measure the *K_d_* between Ync13 and Rga7. (Top) Curve fit showing Rga7 bound fraction versus MBP-Ync13-6xHis concentration on the beads. (Bottom) Coomassie-stained gel of the supernatants showing Rga7-6xHis depletion. Numbers above each lane indicate total MBP-Ync13-6xHis concentration on the beads.

Next we tested if Rga7 and Ync13 directly interact by in vitro binding assays using purified recombinant full length Ync13 and Rga7. MBP-Ync13-6His bound Rga7-6His with a dissociation constant (K_d_) of 10.0 µM (Fig. 2, D and E). These data indicate that Ync13 and Rga7 directly interact. Thus, we conclude that Rng10, Rga7, and Ync13 form a protein complex, which can recruit the glucan synthases Bgs4 (and Smi1) and Ags1 to the plasma membrane at the division plane for secondary septum formation.

### Bgs4 recruitment and distribution at the division plane depends on Rng10, Rga7, and Ync13

Next we asked the functional significance of the detected physical interactions among the proteins in late cytokinesis. We first tested their localization interdependence. We started with Rng10, Rga7, and Ync13 because they colocalized perfectly at cell tips and the division site on the plasma membrane from anaphase until daughter-cell separation (Fig. 1 A and data not shown). Compared to wild type (WT), Ync13 levels at the division site measured by fluorescence intensity were significantly reduced (>85%) in both *rga7Δ* and *rng10Δ* cells (Fig. 3, A and B), although the global Ync13 level was not obviously affected (Fig. 3 C). We next examined the localizations of Rga7 and Rng10 in *ync13Δ* cells. The total Rga7 amount at the division site increased in *ync13Δ* cells, although Rga7 global protein levels were not obviously affected in either *ync13Δ* or *3nmt1-ync13* cells in Western blotting (Fig. 3, D and E). Both Rga7 and Rng10 were more concentrated at the center of division plane in *ync13Δ* cells after the constriction of the contractile ring marked with Rlc1 than in WT cells (Fig. 3, E-G). Interestingly, previous work showed that *ync13Δ* leads to similar central accumulation of the glucan synthase Bgs4 at the division site (Zhu et al., 2018), which resembles the aberrant distribution of Rga7 and Rng10 in *ync13Δ* cells (Fig. 3, E-G).

**Figure 3.**
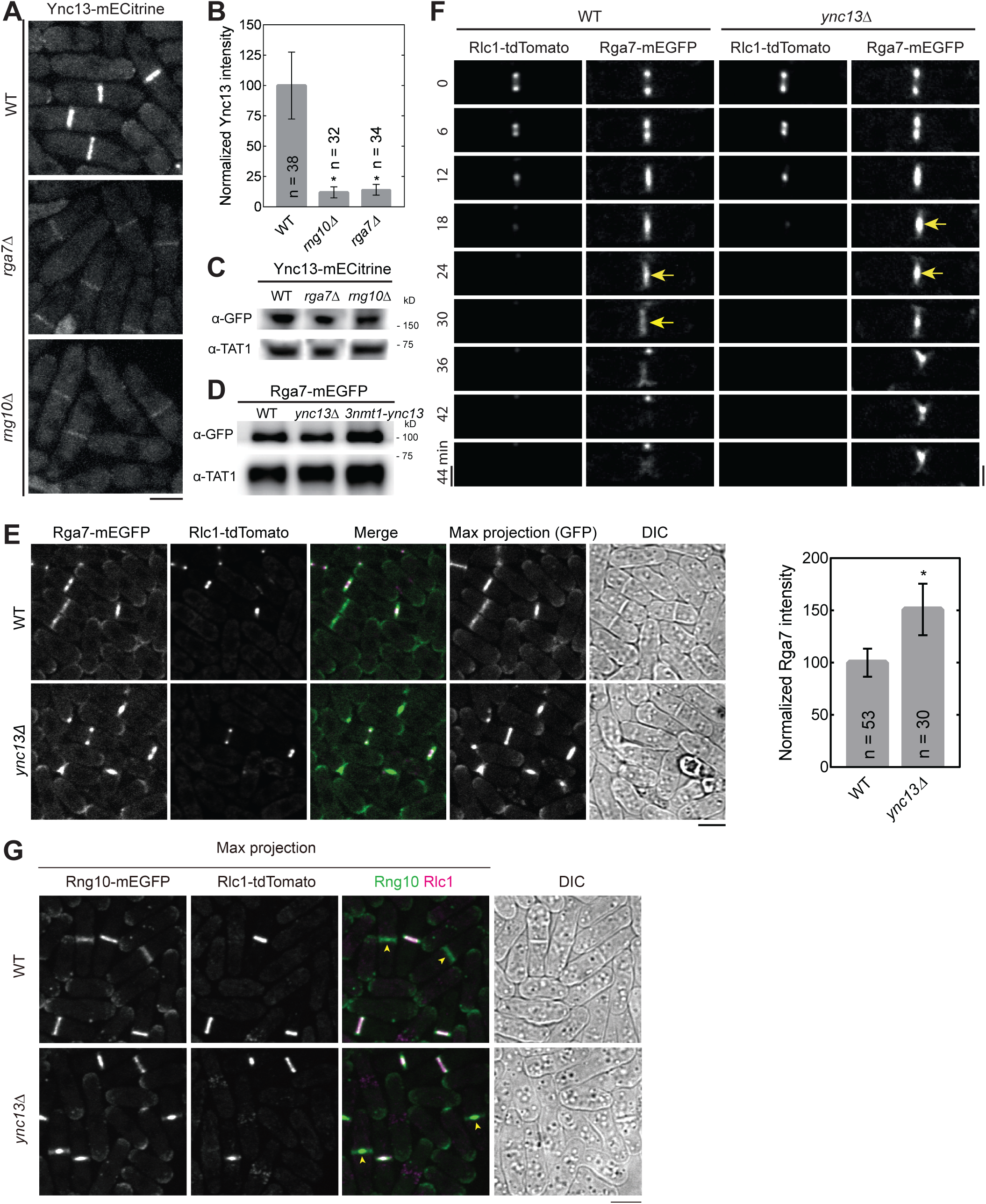
Interdependence of Ync13, Rga7, and Rng10 localization to the division site. **(A and B)** Rng10 and Rga7 are important for Ync13 localization. Micrographs **(A)** and fluorescence intensity at the division site **(B)** of Ync13 in WT, *rga7Δ*, and *rng10Δ*. *p < 0.0001 compared with WT. **(C and D)** Western blotting showing Ync13 **(C)** and Rga7 **(D)** protein levels in cells extracts from the indicated strains. α-TAT1 against tubulin was used as a loading control. **(E and F)** Micrographs and Rga7 intensity at the division site **(E)** and time courses **(F)** showing Rga7 concentration to the center of the division site in *ync13Δ* cells during late cytokinesis. Middle focal plane showing Rga7 localization relative to the contractile ring marked with Rlc1 except the Max projection of GFP in (E). Arrows in (F) mark Rga7 localization at the center of the division plane after ring constriction. **(G)** Micrographs showing Rng10 concentrated to the center of the division site in *ync13Δ* cells during late cytokinesis. Max projection and DIC are shown. Cells were grown exponentially in EMMS liquid media at 25°C for 48 h in (E-G). Bars, 5 μm.

Glucan synthases are essential for maintaining septum thickness and integrity during cytokinesis. Cells of *ync13Δ*, *rga7Δ*, *rng10Δ, ags1*, and *bgs4* mutants all lyse to different degrees during daughter-cell separation dependent on the growth conditions (Cortes et al., 2005; Cortes et al., 2012; Liu et al., 2016; Liu et al., 2019; Zhu et al., 2018). Because the glucan synthases Ags1 and Bgs4 display lower concentrations at the division site in *rga7Δ* and *rng10Δ* cells (Liu et al., 2016; Liu et al., 2019), and Rga7 and Rng10 can mistarget Bgs4 and Ags1 (Fig. 1, D and E), we examined the relationship between mislocalized Rga7, Rng10, Bgs4, and Ags1 in *ync13* mutants. As previously reported (Zhu et al., 2018), compared to WT cells, in *ync13Δ* cells, Ags1 and Bgs4 were more concentrated at cell center during septum maturation (from the end of ring constriction until daughter-cell separation), but Bgs1 was less affected (Fig. 4, A-C). We also found that in *ync13Δ* cells, the intensities of both Ags1 and Bgs4 at the rim of the septum were much lower than in WT after ring constriction, with Bgs4 being more significantly affected compared to Ags1 (Fig. 4, A and B). In contrast, the distribution of Bgs1 was smoother and the intensity at the edge of the division site in *ync13Δ* cell was similar to WT (Fig. 4 C). These were confirmed by the calculated Full Width at Half Maximum (FWHM) from Gaussian fits of Bgs4 and Bgs1 fluorescence intensity across the division site (Fig. 4, D and E). The lack of Ags1 and Bgs4 at the rim of the division plane may influence the rigidity of secondary septum to counter the internal turgor pressure and cause the daughter cell lysis during cell separation. Next we tested if Bgs4 was affected by vesicle tethering and fusion machinery. Consistent with the mistargeting results (Fig. 1 C and Fig. S2 B), Bgs4 levels at the division site decreased in *sec1-M2* and *trs120-ts1* mutants after ring constriction (Fig. S4, A and B).

**Figure 4.**
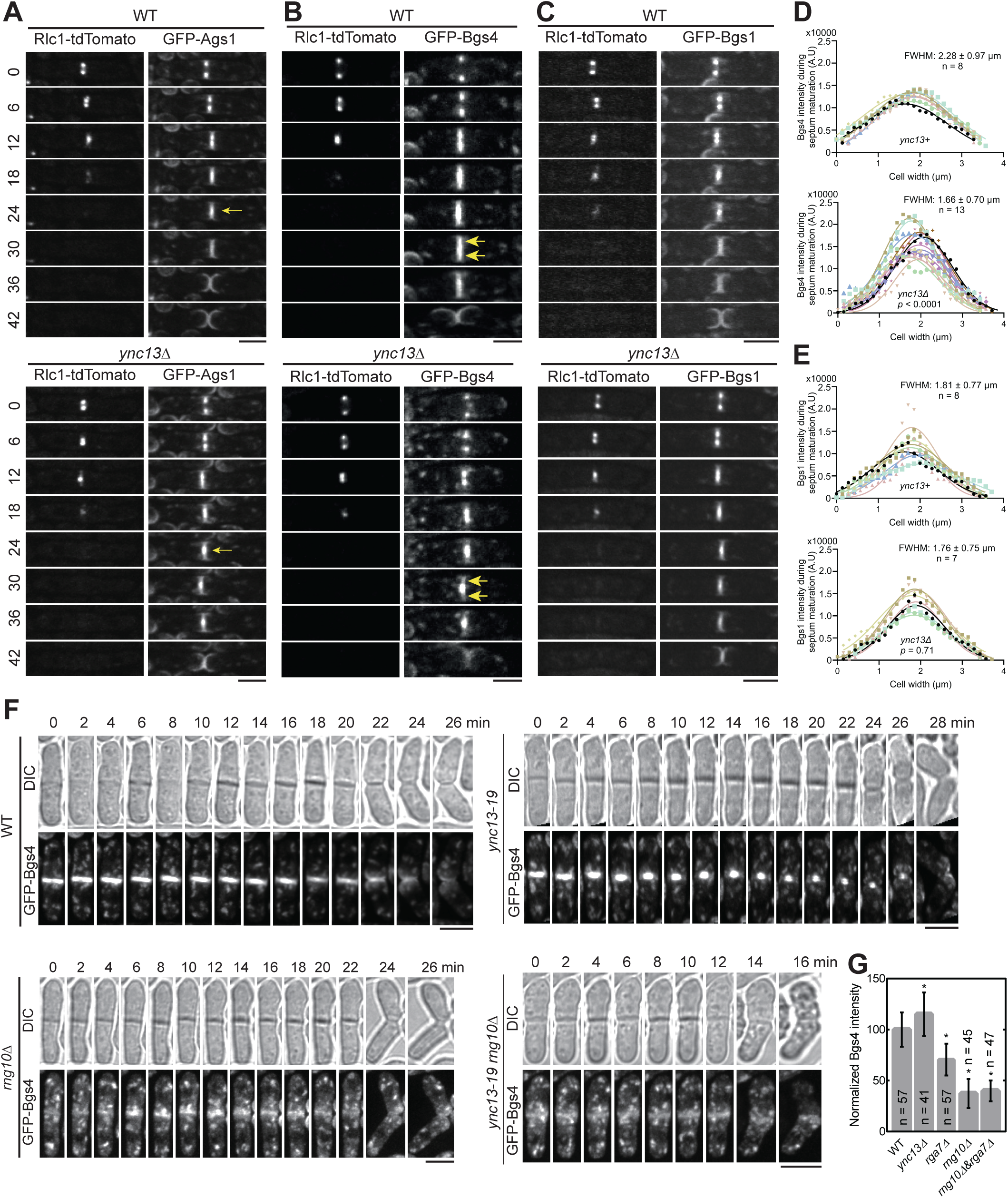
The spatial distribution of Bgs4 in *ync13Δ* cells depends on Rng10. Cells were grown exponentially in EMMS liquid media at 25°C for 48 h before imaging. **(A-C)** Time courses (in min) of Ags1 **(A)**, Bgs4 **(B)**, Bgs1**(C)** distribution at middle focal plane along the division plane relative to the contractile ring marked with Rlc1. Arrows in (A) mark Ags1 concentrated at the center of the division plane after ring constriction and in (B) mark decreased Bgs4 level at the septal edges. **(D** and **E)** Gaussian fits of fluorescence intensity of Bgs4 **(D)** and Bgs1 **(E)** along the division site during septum maturation (after Rlc1 ring disappearance until cell separation) in WT or *ync13Δ* cells. FWHM: mean ± standard deviation. **(F)** Time courses of Bgs4 localization in WT, *ync13-19*, *rng10Δ*, and *rng10Δ ync13-19* cells before and during cell separation. Both daughter cells lysed after cell separation in *rng10Δ ync13-19.* Cells were grown exponentially at 25°C and then shifted to 36°C for 2 h before imaging. Bars, 5 μm. **(G)** Normalized Bgs4 fluorescence intensity at the division site during septum maturation in the indicated strains. *p ≤ 0.0002 compared with WT.

We hypothesize that the *ync13Δ* mislocalizes Rga7 and Rng10, which in turn impacts the distribution of Bgs4 and Ags1 at division site. To test this hypothesis, we first tested Rng10 and Rga7 localization in Bgs4 mutants. Neither Rga7 nor Rng10 showed defects in *bgs4* temperature-sensitive mutants *cwg1-1* and *cwg1-2* (Fig. S4, C and D), suggesting that Bgs4 is not important for the localizations of Rga7 and Rng10. Then we compared Bgs4 localization and intensity in WT, *ync13-19*, *rng10Δ*, and *rng10Δ ync13-19* cells at 36°C. *rga7Δ ync13-19* could not be tested because it was inviable even at 25°C, which is the permissive temperature for *ync13-19* (Zhu et al., 2018). The time-lapse movies showed that the Bgs4 intensity at division site was significantly reduced in *rng10Δ* and *rng10Δ ync13-19* cells, and Bgs4 did not accumulate in the center of the division plane compared to *ync13-19* cells (Fig. 4 F).

Consistently, Bgs4 levels at the division site were significantly lower in *rng10Δ, rga7Δ,* and *rng10Δ rga7Δ* cells (Fig. 4 G). These data indicate Bgs4 is confined at the center of the division plane in *ync13* mutant cells by Rng10 (and Rga7). Thus, Rng10, Rga7, and Ync13 are essential for Bgs4’s localization, recruitment, and normal spatial distribution at the division site.

### Both Rga7 and Rng10 are involved in TRAPP-II complex dependent exocytosis at the division site

Both *rga7Δ* and *rng10Δ* cells show compromised recruitment of glucan synthases at division site and have defective septum (Arasada and Pollard, 2015; Liu et al., 2016; Liu et al., 2019; Martin-Garcia et al., 2014). *rga7Δ rng10Δ* double deletion cells are inviable in rich media at 25°C without osmotic stabilizer sorbitol (Liu et al., 2016). Next, we examined how Rga7 and Rng10 affect the trafficking of the glucan synthase containing vesicles, focusing on the TRAPP-II complex because both Rga7 and Rng10 can mistargeting Trs120 to mitochondria. The TRAPP-II complex promotes vesicle tethering for exocytosis along the cleavage furrow and its subunit Trs120, which is sufficient for TRAPP-II localization, co-localizes with Bgs4 containing vesicles (Cai et al., 2005; Choi et al., 2011; Wang et al., 2016). Trs120 can mistarget Bgs4 to mitochondria and dynamically localizes to the division site during and after ring constriction (Wang et al., 2016). In *rga7Δ*, *rng10Δ*, and *rga7Δ rng10Δ* (shifting from medium with sorbitol to the one without sorbitol) cells, we found that Trs120 intensity at division site decreased significantly, while it increased in *ync13Δ* cells (Fig. 5, A and B; arrowheads). Trs120 was also more concentrated at the center of the division plane in *ync13Δ* cells (Fig. 5 A). The accumulation of Trs120 outside of the division site in cells with a nearly closed or closed septa was detectable in the sum projection images of 2-min continuous movies at middle focal plane in *rga7Δ*, *rng10Δ*, and *rga7Δ rng10Δ* but not in WT cells expressing Trs120-3GFP, indicating membrane tethering defect (Fig. S5, arrowheads).

**Figure 5.**
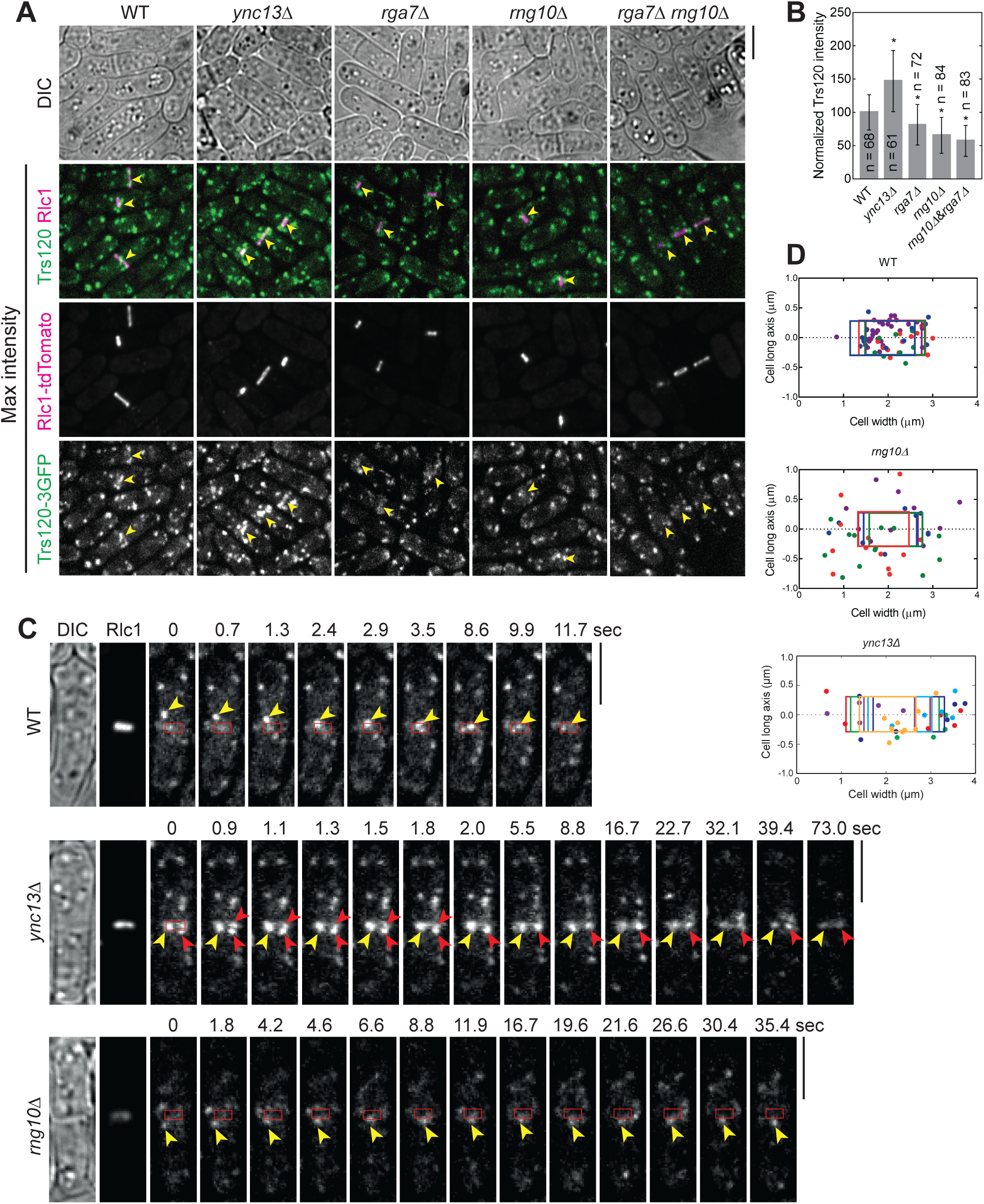
The Rga7-Rng10-Ync13 module regulates TRAPP-II complex-mediated vesicle tethering and fusion at the division site. Cells were grown exponentially in EMMS liquid media at 25°C for ∼48 h before imaging. **(A)** Trs120-3GFP localization in WT and *ync13Δ*, *rga7Δ*, *rng10Δ*, and *rga7Δ rng10Δ* cells. Arrowheads mark the Trs120 at division site in cells with constricting ring labeled with Rlc1-tdTomato. **(B)** Quantification of Trs120 intensity at division site in cells with a constricting ring as in (A). *p < 0.0001 compared with WT. **(C)** Trs120 puncta (arrowheads) move to the division site during ring constriction in WT, *rng10Δ*, and *ync13Δ* cells. Red boxes mark the ring position. Please also see the Videos 1-3. **(D)** Distribution of final tractable docking sites (each cell with colored dots that match the box color) of Trs120 labeled puncta in 2-min movies during ring constriction. The color boxes mark the ring position as shown in (C). Bars, 5 μm.

We also tracked Trs120 vesicle movement during the late stage of ring constriction. In WT cells, Trs120 puncta were delivered to the leading edge of the cleavage furrow mainly concentrated within the region close to the ring (Fig. 5, C and D; Video 1). However, in *rng10Δ* cells, Trs120 puncta rarely docked near the leading edge of the division site, which may be due to the delayed vesicle tethering (Fig. 5, C and D; Video 2). Interestingly, in *ync13Δ* cells, more Trs120 concentrated within the division plane during and after ring constriction than in WT cells (Fig. 5, A, C, D; Fig. S5; Video 3). Consistently, in both *rng10Δ* and *ync13Δ* cells, it took longer for vesicles to fuse with the plasma membrane at division site compared to WT (Fig. 5 C). Thus, Rga7 and Rng10 participate in vesicle tethering mediated by the TRAPP-II complex for septum formation.

### Electron microscopy reveals defective septum in *ync13Δ* cells

Because many more *ync13Δ* cells lyse when grown in YE5S rich medium than in the EMM5S minimal medium, we observed septum morphology after shifting cells from YE5S with sorbitol to YE5S by electron microscopy. After growing *ync13Δ* cells in YE5S for 3.5 h, >50% forming or closed septa were defective (Fig. 6, A-C). This figure is likely underestimated because many cells had lysed before the high pressure freezing to preapare the samples for electron microscopy. Compared to WT, the septa in *ync13Δ* cells are distorted, curved, wavy, uneven, and/or thinner (Fig. 6, A and B), suggesting improper septum formation, especially the secondary septum. The septum in many cells was thin or missing at the edge of the division plane during daughter-cell separation, which led to cell lysis (Fig. 6 B, two cells at the bottom right). The primary septum had no obvious defects in *ync13Δ* cells (Fig. 6 B, arrowheads). We also observed some electron dense filamentous materials at the leading edge of the forming septum in some *ync13Δ* cells (arrow), which was rare in WT cells. These defects occured earlier during septum formation and were more severe than those described in published data when cells were grown in the EMM5S media (Zhu et al., 2018). The electron microscopy results support the idea that Ync13 is important for recruitment and maintenance of glucan synthases Bgs4 and Ags1 on the plasma membrane for secondary septum formation during cytokinesis.

**Figure 6.**
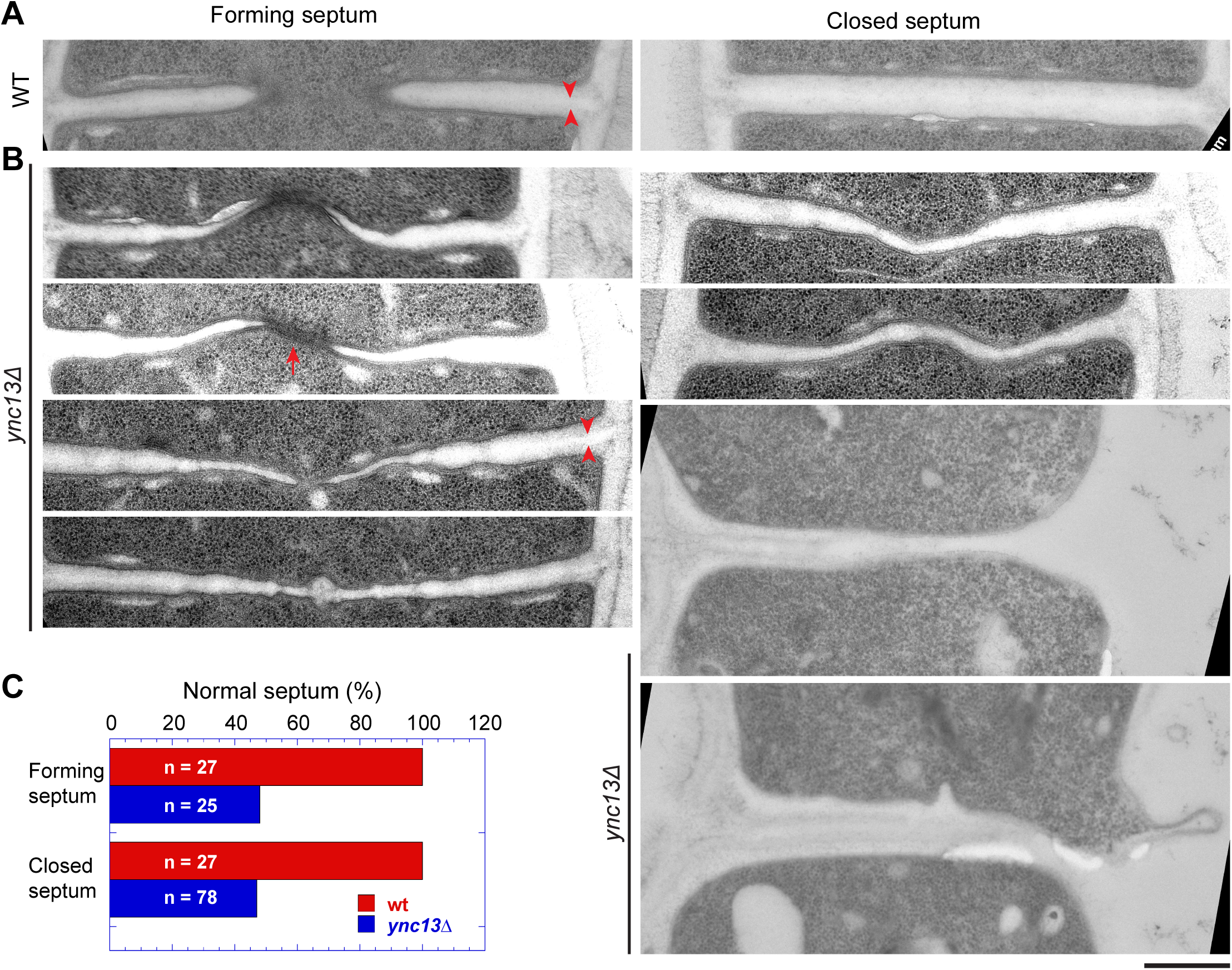
The septum is defective in *ync13Δ* cells revealed by electron microscopy. Electron microscopy thin sections were rotated and interpolated (bicubic) using Fiji so the septum is horizontal. Then the septum region was cropped and shown. Left: forming septum; Right: closed septum or daughter-cell separation and cell lysis (from the right side, the bottom two cells). Cells were grown exponentially at 25°C in YE5S + 1 M sorbitol for ∼48 h and then washed into YE5S without sorbitol and grown for 3.5 h before collection for high pressure freezing. Arrowheads, mark examples of the primary septum. Arrow, marks an example of the electron dense materials near the leading edge of the septum. **(A)** *ync13^+^* (WT) cells. **(B)** Representative *ync13Δ* cells with abnormal septa. **(C)** Quantification of percentage normal septa in *ync13^+^* (WT) and *ync13Δ* cells. The septa in (B) are defined as abnormal. Bar, 500 nm.

## Discussion

The interplay between membrane trafficking and septum formation or extracellular matrix remodeling is critical for maintaining cell integrity and successful cytokinesis from yeast to mammalian cells (Balasubramanian et al., 2004; Izumikawa et al., 2010; Mizuguchi et al., 2003; Nishihama et al., 2009; Xu and Vogel, 2011a; Xu and Vogel, 2011b). Our findings elucidate the central roles of the Ync13-Rga7-Rng10 complex in coordinating selective vesicle tethering, docking, and fusion mediated by the TRAPP-II complex and SM protein Sec1 at the cleavage furrow (Fig. 7). This module ensures precise and timely secondary septum formation by Bgs4 and Ags1 and prevents cell lysis during daughter-cell separation (Fig. 7). Bgs1, recruited by Sbg1 and the contractile ring anchored by the anillin Mid1 and F-BAR protein Cdc15, is responsible for the primary septum formation (Arasada and Pollard, 2014; Arellano et al., 1997; Bhattacharjee et al., 2023; Cortes et al., 2002; Davidson et al., 2016; Garcia et al., 2006a; Nakano et al., 1997; Ren et al., 2015; Roberts-Galbraith et al., 2010; Sethi et al., 2016; Wang et al., 2016); while septins and the exocyst are mainly concentrated at the rim of the division plane to recruit the glucanase Eng1 to digest the primary septum during daughter-cell separation (Alonso-Nunez et al., 2005; Berlin et al., 2003; Martin-Cuadrado et al., 2005; Pérez et al., 2015; Santos et al., 2005; Singh et al., 2024; Tasto et al., 2003; Wang et al., 2003; Wang et al., 2015) (Fig. 7).

**Figure 7.**
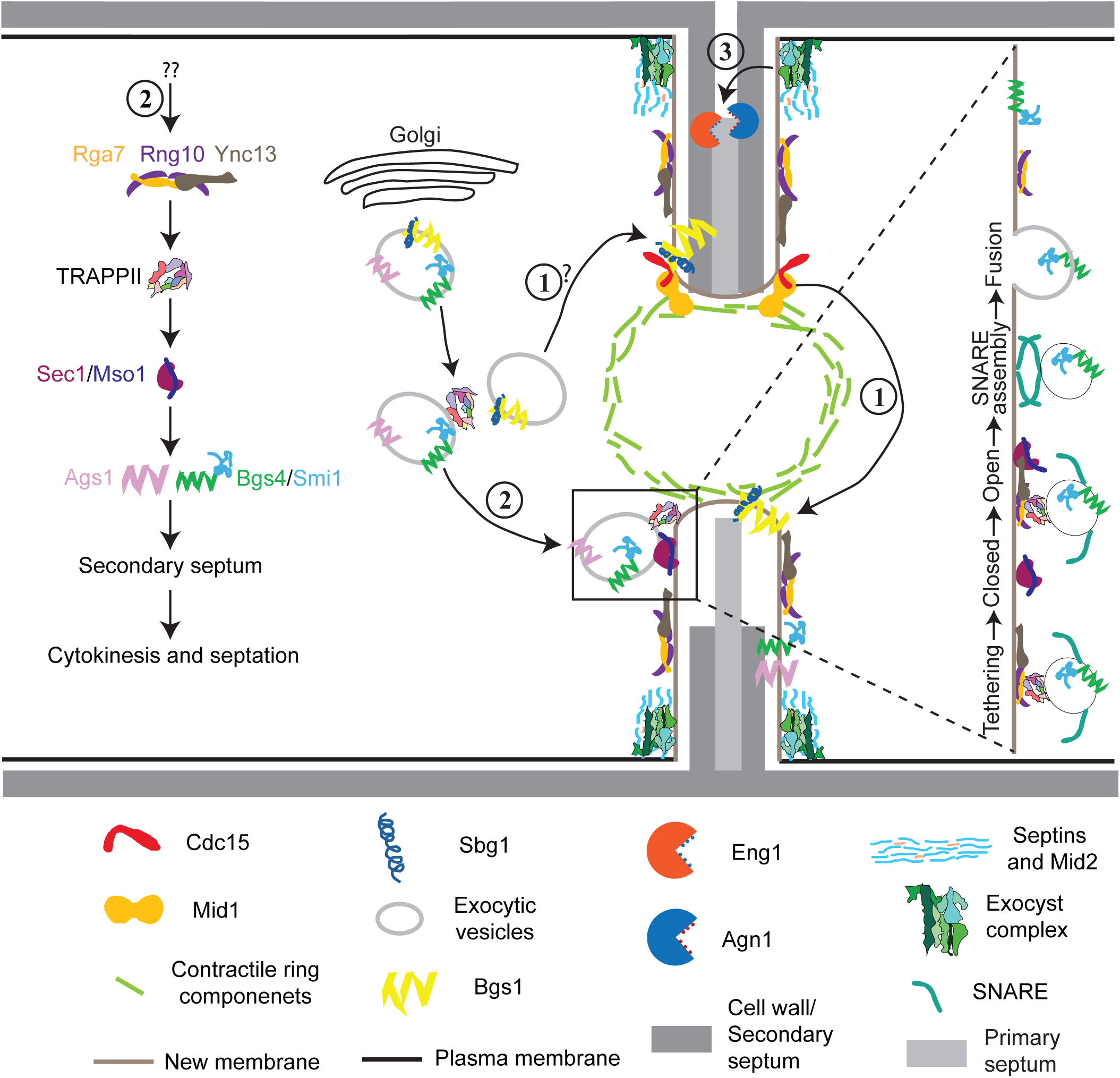
Working model for the major pathways of plasma membrane deposition and septum formation during cytokinesis. The septum is illustrated asymmetrically intentionally. Top half septum shows digestion of the primary septum; bottom half septum shows septum formation. (1) The anillin Mid1 and F-BAR protein Cdc15 anchor the contractile ring to the plasma membrane, which recruits the β(1,3)-glucan synthase Bgs1 and its adaptor Sbg1 for primary septum formation. (2) Association of the F-BAR protein Rga7 and coiled-coil protein Rng10 with the Munc13/Unc13 protein Ync13 recruits the TRAPP-II complex along the cleavage furrow, which tethers vesicles and primes the SM protein Sec1/Mso1 for the SNARE complex assembly to trigger vesicle fusion with the plasma membrane independent of the exocyst complex. This pathway maintains normal levels and distribution of gluacan synthases Bgs4/Smi1 and Ags1 for secondary septum formation. (3) The anillin Mid2 and septin rings restrict most of the exocyst complex to the rim of cleavage furrow to tether secretory vesicles loaded with glucanases Eng1 and Agn1 and other cargos for digestion of primary septum to trigger daughter-cell separation.

### Rga7, Rng10, and Ync13 cooperatively regulate vesicle tethering and fusion during cytokinesis

Our findings have several mechanistic implications for how vesicle tethering and fusion are coupled during cytokinesis. First, the interaction of Ync13 with Sec1 suggests that Ync13 functions as a SNARE-priming factor at the division site before SNARE complex assembly. SM protein Sec1 and its binding partner Mso1 bind syntaxin-family SNAREs and are essential for polarized secretion (Carr et al., 1999; Castillo-Flores et al., 2005; Gerien et al., 2020; Weber-Boyvat et al., 2011). In other systems, SM proteins work in tandem with Munc13 proteins to assemble fusogenic SNARE complexes (Bhaskar et al., 2024; Khodthong et al., 2011; Ma et al., 2011; Shu et al., 2020). The fact that mistargeted Ync13 recruits Sec1 to mitochondria suggests a direct role for Ync13 in vesicle fusion, analogous to the established function of mammalian Munc13 proteins (Guan et al., 2008; Khodthong et al., 2011; Ma et al., 2011; Shu et al., 2020), which need to be tested by in vitro reconstitution assays in future studies. Second, Ync13, by binding to the lipid binding protein complex Rga7/Rng10 (Liu et al., 2016; Liu et al., 2019), is well positioned to help tether or recruit vesicles to the division site, where its interaction with Sec1 could facilitate the transition from a tethered vesicle to a docked, SNARE-engaged vesicle ready for fusion. Third, Ync13’s direct interaction with the Rga7/Rng10 complex links it indirectly to the TRAPP-II vesicle tethering complex. We observed that Rga7 and Rng10 selectively interact with the TRAPP-II but not with the exocyst complex in mislocalization experiments. The absence of interaction of Ync13, Rng10, or Rga7 with the exocyst complex indicates that Ync13 performs a TRAPP-II specific function on vesicle tethering. This is reinforced by the distinct accumulation of Trs120 puncta in *rng10Δ* and *ync13Δ* mutants. In *rng10Δ* cells, TRAPP-II–marked vesicles often fail to arrive at the cleavage furrow and accumulate outside the division site. In contrast, in *ync13Δ* cells, vesicles arrive at the division furrow but linger there longer before fusion. We speculate that, similar to Munc13 (Dulubova et al., 2005; Leitz et al., 2024; Lu et al., 2006), the function of Ync13 may be attributed to a post vesicle docking stage after tethering. When the TRAPP-II-mediated tethering is compromised by *rga7Δ* and *rng10Δ*, the vesicles cannot efficiently contact with its target membrane surface and move outside of the division site. However, in *ync13Δ* cells, the physical interaction between vesicles and the target membrane may have been established after tethering process, but the docking process is affected, which leads to the delayed vesicle fusion. Roles of the TRAPP-II complex in cytokinesis have been reported in *Drosophila* and plant cells (Robinett et al., 2009; Rybak et al., 2014). Taken together, we propose that the Rga7/Rng10 complex serves as a scaffold for vesicle tethering by the TRAPP-II complex and interacts with Ync13 to facilitate the assembly of the SNARE complex and promote membrane fusion.

Munc13 functions in neurotransmitter release by priming vesicle tethering process for sudden release of synaptic vesicle pool (Ma et al., 2013). This ensures a reliable and speedy neurotransmission following synaptic collapse. Ync13 regulates both exocytosis and endocytosis. However, the exact mechanisms and its interactions with other proteins were unknown before this study. Our previous studies show that Rga7 binds the C-terminal portion of Rng10 to increase Rga7’s avidity with the plasma membrane (Liu et al., 2016; Liu et al., 2019). Our studies here indicate that Rga7/Rng10 interacts with Ync13, which interacts with the SM protein complex Sec1/Mso1. These proteins play a crucial role for tethering the secretory vesicles by regulating the TRAPP-II and SNARE complexes, which are essential for fusion of vesicles with target membranes (Fig. 7).

### Selective recruitment of the glucan synthases for the secondary septum by the Rga7-Rng10-Ync13 module at the division site

The transmembrane glucan synthases Bgs1, Bgs4, and Ags1 are essential for septum formation during cytokinesis and are maintained at the proper levels on the plasma membrane via membrane trafficking (Cortes et al., 2005; Cortes et al., 2002; Cortes et al., 2007; Cortés et al., 2015; Cortes et al., 2012; Ishiguro et al., 1997; Katayama et al., 1999; Liu et al., 1999). The primary septum in fission yeast is composed of mainly linear β-1,3 glucan chains synthesized by Bgs1 that plays similar role as chitin in budding yeast (Cabib et al., 1993). Bgs1 concentrates behind the contractile ring, while Ags1 and Bgs4 localize at the cleavage furrow on both layers of the new plasma membrane to construct secondary septum. At low restrictive temperatures, *sid2* mutant lethality is partially rescued by upregulating Rho1. Thus, it has been suggested that the SIN activates Rho1, which in turn activates the glucan synthases (Alcaide-Gavilán et al., 2014).

Our data demonstrate that the Rga7-Rng10-Ync13 module specifically recruits Ags1 and Bgs4 for secondary septum assembly, but not Bgs1 for primary septum synthesis. Ectopic targeting of Rga7 or Rng10 is sufficient to redirect vesicles containing Bgs4 and Ags1 to mitochondria. However, Bgs1 is not mislocalized under these conditions. When overexpressed, Ync13 can also pull Bgs4 and Ags1 to mitochondria, presumably through Ync13’s binding to Rga7/Rng10. Importantly, the proper spatial distribution of Bgs4 and Ags1 relies on the functional interaction between Ync13, Rga7, and Rng10. In WT cells, Rga7, Rng10, Ync13, Bgs4, and Ags1 colocalize at the division plane and are mostly evenly distributed around the edges of the invaginating septum and throughout the maturing septum. In contrast, in *ync13Δ* cells, all these proteins abnormally concentrate at the leading edge of the furrow and later at the center of the division plane, forming a flattened convex structure. This mislocalization depends on Rga7/Rng10 and leads to a significant reduction in Bgs4 (and to a lesser extent Ags1) near the septum periphery, highlighting the critical role of Ync13 in maintaining proper enzyme distribution. The lack of proper Ags1/Bgs4 deposition on the plasma membrane at the septum edges likely compromises the thickness and rigidity of the new cell wall, explaining why *ync13Δ*, *rga7Δ*, and *rng10Δ* mutants all undergo cell lysis during cell separation. In contrast, the primary septum synthase Bgs1 remains correctly localized even in *ync13Δ* cells, suggesting that primary septum formation follows a Ync13/Rga7/Rng10-independent pathway. Thus, our study highlights the distinct roles of Rga7–Rng10–Ync13 module in controlling the late-stage cell wall assembly: it specifically regulates the formation of the secondary septum and maintains cell wall integrity by recruiting the secondary septum glucan synthases Bgs4 and Ags1 at the division site.

Our findings have two important mechanistic implications. First, the finding that the Rga7-Rng10-Ync13 complex only recruits Bgs4 and Ags1, but not Bgs1, strongly indicates that there are at least two distinct types of secretory vesicles to transport cell wall synthases. In the budding yeast *S. cerevisiae*, different types of secretory vesicles have been reported (Conibear and Stevens, 1998; Harsay and Bretscher, 1995). The constitutive or low density secretory vesicles deliver plasma membrane proteins, cell wall components, and enzymes necessary for cell growth from trans-golgi network. Invertase-containing or high density secretory vesicles transport stress-induced cargos, such as the enzyme invertase (Ford et al., 2021; Kienzle and von Blume, 2014). Second, the central and leading edge localization of Rga7, Rng10, Bgs4, and Ags1 in *ync13* mutants suggests that the Bgs4 and/or Ags1-containing vesicles are mis-directed to where the Bgs1 vesicles are normally targeted, perhaps using the machinery that the Bgs1 vesicles use. It will be interesting to test these possibilities in future studies.

Rho GTPases are small molecular switches that regulate multiple cellular processes including cytokinesis (Hall, 2012). Of the six Rho GTPases (Rho1-5 and Cdc42) in fission yeast, Rho1 and Rho2 play crucial roles in maintaining cell integrity during cytokinesis (Arellano et al., 1999; Calonge et al., 2000; Garcia et al., 2006b; Perez and Rincon, 2010; Sánchez-Mir et al., 2014). Activated mainly by Rho GEF Rgf3 and its adapter arrestin Art1, GTP-bound Rho1 activates β-glucan synthases Bgs1 and Bgs4 and protein kinase Cs Pck1 and Pck2 (Arellano et al., 1999; Arellano et al., 1996; Davidson et al., 2015; Morrell-Falvey et al., 2005; Mutoh et al., 2005; Ren et al., 2015; Sánchez-Mir et al., 2014; Tajadura et al., 2004). Similarly, Rho2 functions in the cell integrity pathway and activates α-glucan synthase Ags1 for septum formation (Calonge et al., 2000; Sánchez-Mir et al., 2014). However, it is unclear how Rho GTPases regulate recruitment, distribution, and maintenance of glucan synthases.

More studies are available in budding yeast regarding the regulations of septum or cell wall synthases. The primary septum in the budding yeast *S. cerevisiae* consists of the chitin synthesized by chitin synthases Chs1 and Chs2, which functions similarly to the primary septum of fission yeast consisting of linear β(1,3)glucan (Cabib et al., 1993; Oh et al., 2012; Schmidt et al., 2002; Walker et al., 2013; Zhang et al., 2006). Cdc14 dephosphorylates Chs2 for its localization at the septation site from ER (Chin et al., 2012; Zhang et al., 2006). Chs3 is responsible for the formation of a chitin ring at the emerging bud and of the chitin dispersed in the cell wall. Chitin synthase III complex also contains Chs4 that anchors to the plasma membrane and interacts with Chs3 and Bni4 (DeMarini et al., 1997). Septins interact with Bni4 to hold this complex together which help the chitin synthase to synthesize sufficient chitins for the primary septa. Chitin synthases localize in Rho1 dependent manner (Yoshida et al., 2009).

The secondary septum in budding yeast is made up of β-glucans with minor amount of chitin. Rho1 and Rho2 activate FKS1 by interacting with the glycosyltransferase domain and the transmembrane helix (Drgonová et al., 1996; Levin, 2005; Onishi et al., 2013; Qadota et al., 1996). These interactions induce conformational changes to push growing β-1,3-glucan chain (Li et al., 2025). Alternatively secondary septum synthesis can also be achieved by transcriptional activation of *FKS2* and *GFA1* genes by the cell wall integrity pathway (Roncero et al., 2021).

Cdc42 negatively regulates secondary septum formation (Das et al., 2009; Onishi et al., 2013; Rincon et al., 2007; Tatebe et al., 2008). However, it is still poorly understood how the glucan synthases FKS1/2 are recruited to the division site in budding yeast.

By identifying the Rga7-Rng10-Ync13 module as the main pathway for proper recruitment and distribution of the glucan synthases Bgs4 and Ags1 at the division site for secondary septum formation, our current study provides significant insights into the regulatory mechanisms of glucan synthases, which are ideal targets for antifungal drugs. However, cells must possess other unidentified mechanisms for Bgs4 and Ags1 recruitment because *ync13Δ* or *rga7Δ rng10Δ* cells can survive in medium with sorbitol or in minimal medium, although not in rich medium. Rga7 is Rho2 GAP (Arasada and Pollard, 2015; Soto et al., 2010; Villar-Tajadura et al., 2008), thus it is possible that the Rga7-Rng10-Ync13 module regulates Bgs4 and Ags1 via Rho GTPases. However, the functions of Rga7’s GAP domain remain poorly understood (Arasada and Pollard, 2015; Liu et al., 2019; Martin-Garcia et al., 2014). Moreover, no connection between FKS1/2 and the homologs (RGD1 and YOR296W, respectively) of Rga7 and Ync13 in budding yeast have been established.

In summary, in this study we find that Ync13, Rga7, and Rng10 form a regulatory module that links vesicle trafficking with the precise targeting and distribution of glucan synthases Bgs4 and Ags1 to ensure efficient septum formation. Ync13 plays a key role in both vesicle fusion and spatial distribution of Rga7/Rng10, underscoring its crucial function in cytokinesis. Moreover, the selective interactions between Rga7/Rng10, TRAPP-II, and glucan synthases fine-tune vesicle targeting specificity. These findings deepen our understanding of how membrane trafficking processes are spatially and temporally coordinated to enable successful cytokinesis.

## Materials and methods

### Strain construction and growth methods

The strains used in this study are listed in Table S1. Strains were constructed by tagging genes at endogenous chromosomal loci using standard PCR and homologous recombination-based gene targeting method (Bähler et al., 1998) so fusion proteins were expressed under the control of native promoters. The exceptions are tagged glucan synthases Bgs1, Bgs4, and Ags1, which are regulated by the native promoters but integrated at the leu1 loci with the endogenous copies deleted (Cortes et al., 2005; Cortes et al., 2002; Cortes et al., 2007; Cortes et al., 2012; Hochstenbach et al., 1998; Munoz et al., 2013); and the *3nm1-ync13* strains. Cells were grown at exponential phase (OD595 < 0.5) at 25°C in YE5S (yeast extract with five supplements) liquid medium for ∼48 h before imaging (more details below) or other experiments except where noted.

To tag Ync13 with 3xFLAG, the fragment of GGSGGS-3xFLAG was first inserted at the SmaI site in pREP3x plasmid vector. Full-length (FL) Ync13 cDNA was cloned into pREP3x-GGSGGS-3xFLAG between the *3nmt1* promoter and GGSGGS-3xFLAG tag by Gibson assembly (Gibson et al., 2009). The plasmids pREP3x-Ync13-FL-3xFLAG and pREP3x-3xFLAG (control) were then transformed into protease deficient strain TP150 *leu1-32 SM902* (Lord and Pollard, 2004; Ti and Pollard, 2011) and positive transformants were selected on EMM5S - leucine medium. The positive colonies were grown on EMM5S - leucine liquid medium to OD595 = 1.0 before collection and lyophilization.

### Mass spectrometry

We used the previously described protocol for purification of S-tagged proteins with several modifications to identify the Ync13 binding partners (Liu et al., 2010). Approximately 2 g of lyophilized cells (TP150 strains with plasmid pREP3x-Ync13-FL-3xFLAG or pREP3x-3xFLAG control) were broken by grinding with a mortar and pestle at room temperature until the cells became a homogenous fine powder. For protein extraction, the cell powders were thoroughly mixed with 25 ml of cold HK extraction buffer (25 mM Tris, pH 7.5, 0.5% NP-40, 300 mM NaCl, 5 mM EDTA, 15 mM EGTA, 60 mM β-Glycerophosphate, 500 µM Na_3_VO_4_, 10 mM NaF, 1 mM PMSF, 1 mM DTT, and protease inhibitors [Roche]). The cell extracts were cleared by two rounds of centrifugation at 4°C (21,000 rpm for 30 min, 21,000 rpm for 10 min). Then cell extracts were incubated with 200 µL anti-FLAG M2 affinity gel (A2220, Millipore) at 4°C for 2 h. The beads were collected by centrifugation at 4,000 rpm and washed once with an equal volume of HK extraction buffer, 4x with an equal amount of washing buffer (25 mM Tris, pH 7.5, 300 mM NaCl, 5 mM EDTA, 500 µM Na_3_VO_4_, 10 mM NaF, 1 mM PMSF, and 1 mM DTT), and 2x with 1 ml of washing buffer. The proteins on the beads were eluted by incubating with 500 µl of 200 μg/ml 3xFLAG peptide (F4799, Sigma-Aldrich) at 4°C for 30 min. For mass spectrometry analysis, the samples were run through the SDS-PAGE gel. Protein bands except the Ync13 band were excised as one sample and processed for mass spectrometry (Mass Spectrometry and Proteomics Facility, The Ohio State University).

### Confocal microscopy and image analysis

Cells from −80°C stocks were streaked onto YE5S plates and grown at 25°C for about 2 d and then fresh cells were inoculated into YE5S liquid medium and grown in the log phase for ∼48 h at 25°C before imaging except where noted. For strains requiring osmotic stabilizer to survive, such as strains with *ync13Δ, ync13-19 rng10Δ*, *rng10Δ rga7Δ*, and some strains with Tom20-GBP, cells were woken up on YE5S + 1.2 M sorbitol plate and grown in YE5S + 1.2 M sorbitol liquid medium at log phase for 36 to 48 h, and then were washed with YE5S without sorbitol and grown in YE5S for 4 h before imaging except where noted. For comparison, some strains with *ync13Δ, ync13-19, rga7Δ,* and *rng10Δ*, and their double mutants of necessary combinations were grown in EMM5S for 48 h before imaging. These strains have less severe phenotype or cell lysis in EMM5S than in YE5S rich medium.

Confocal microscopy was done as previously described (Davidson et al., 2015; Gerien et al., 2020; Longo et al., 2022; Wang et al., 2015). Briefly, cells were collected by centrifugation at 3,000 rpm for 30 s and washed once with 1 ml EMM5S and once 1 ml EMM5S containing n-propyl gallate at a final concentration of 5 µM to reduce autofluorescence and protect cells from free radicals during imaging (Giloh and Sedat, 1982; Laporte et al., 2011). We imaged cells on glass slides with a gelatin pad (20% gelatin in EMM5S + 5 µM n-propyl gallate) at ∼23°C. For long movies, cells were washed with 1 ml EMM5S + 5 µM n-propyl gallate and placed onto a coverglass-bottom dish (Delta TPG Dish; Biotechs, Butler, PA, United States). For fluorescence microscopy at 36°C, the cells were grown at 25°C for ∼2 d and then shifted to 36°C and grown for a given time (see figure legends). Before imaging, cells were washed and concentrated in pre-warmed YE5S liquid medium with 5 µM n-PG. Then 10 μl of the concentrated cells were spotted onto a coverglass-bottom dish, covered with the prewarmed YE5S agar, and imaged at 36°C in a preheated climate chamber (stage top incubator INUB-PPZI2-F1 equipped with UNIV2-D35 dish holder; Tokai Hit, Shizuoka-ken, Japan).

For most fluorescence images and time-lapse movies, cells were imaged using a spinning-disk confocal system (UltraVIEW Vox CSUX1 system; PerkinElmer, Waltham, MA) with 440-, 488-, 515-, and 561-nm solid-state lasers and a back thinned electron-multiplying charge-coupled device (EMCCD) camera (C9100-23B; Hamamatsu Photonics, Bridgewater, NJ) on a Nikon Ti-E microscope without binning (Davidson et al., 2016; Singh et al., 2024). For Fig. 1F, cells were imaged using a Nikon spinning-disk confocal system (W1 + SoRa) with 488 and 561-nm solid-state lasers and an ORCA-Quest qCMOS camera (C15550; Hamamatsu Photonics, Bridgewater, NJ) on a Nikon Eclipse Ti-2E microscope with 2×2 binning (Ye et al., 2025).

Images were analyzed using Volocity (PerkinElmer) and ImageJ/Fiji (National Institutes of Health, Bethesda, MD). Fluorescence images shown are single middle focal plane or maximum-intensity projections of image stacks with 0.5 µm spacing. To measure protein levels across the division plane, the cells after constriction of the contractile ring (Rlc1 ring constricted to a spot at cell center) were chosen and rotated so that the septa were horizontal. A 23 × 6 pixel region of interest (ROI) was drawn to cover the protein signal at division site. The plot profile of the ROIs was recorded.

We tracked secretory vesicles similarly as before (Wang et al., 2016; Zhu et al., 2018). Briefly, the middle focal plane of cells was imaged with a speed of 2–5 frames per second (fps) for the channel with fluorescently labeled vesicles in 2-min movies. The Rlc1 channel was imaged once every minute. DIC images were taken as snapshots immediately before and after the fluorescence movies to make sure no focal shifting. The movements of vesicles were tracked manually using ImageJ plug-in mTrackJ (Meijering et al., 2012). The data coordinates were then transformed by Matlab software so that the septa were horizontal. The cell width was normalized to 4 µm before plotting. Statistical analyses were performed using two tailed Student’s t test in this study.

### Plasmid construction, protein purification, in vitro binding assays

The fragment of MBP-TEV-GGSGGS was first cloned into pET21a vector before BamHI site by Gibson assembly to construct pET21a-MBP vector (Gibson et al., 2009). Ync13 FL cDNA was cloned into pET21a-MBP vector between the GGSGGS linker and the 6His tag by Gibson assembly. Full length Rga7 was cloned into the pET21a vector between the T7 tag and the 6His tag by Gibson assembly. The constructs were confirmed by sequencing.

We purified recombinant proteins by transforming the plasmids into BL21 (DE3) pLysS cells (Novagen). MBP-Ync13-6xHis expression was induced with 0.2 mM IPTG at 17°C for 36-48 h. Rga7-6xHis was expressed with 0.5 mM IPTG at 25°C for 15 h. Purifications of 6His-tagged proteins were carried out as previously described (Wu et al., 2008; Zhu et al., 2013).

Briefly, the proteins were purified with Talon metal affinity resin (635501; Clontech, Mountain View, CA) in extraction buffer (50 mM sodium phosphate, pH 8.0, 400 mM NaCl, 10 mM *β*-mercaptoethanol, 1 mM PMSF, and 20 mM imidazole) with EDTA-free protease inhibitor tablet (Roche) and eluted with elution buffer (50 mM sodium phosphate, pH 8.0, 400 mM NaCl, 10 mM *β*-mercaptoethanol, 1 mM PMSF, and 300 mM imidazole). The purified proteins were then dialyzed into the binding buffer (137 mM NaCl, 2 mM KCl, 10 mM Na_2_HPO_4_, 2 mM KH_2_PO_4_, 0.5 mM dithiothreitol, and 10% glycerol, pH 7.4).

For in vitro binding assays between MBP-Ync13-6xHis and Rga7-6xHis, purified proteins were dialyzed into the binding buffer. We incubated 1 ml MBP-Ync13-6His (2 µM) or 75 µl MBP-6xHis (27 µM) control with 500 µl Amylose beads for 1 h at 4°C and washed the beads 8x with 1 ml of the binding buffer each time to remove unbound proteins. Then 1 ml Rga7-6xHis (10 µM) was incubated with the 100 µl beads with MBP-Ync13-6xHis or MBP-6xHis for 1 h at 4°C. After washing 4x with 1 ml of the binding buffer each time, the beads were boiled with sample buffer for 5 min. Then the samples were run on SDS–PAGE gel and detected with Coomassie Blue staining. For measuring the Kd between MBP-Ync13-6His and Rga7-6His, we followed the methods and guidelines as described (Lee et al., 1999; Liu et al., 2019; Pollard, 2010). Rga7-6xHis at 4 µM was titrated with MBP-Ync13-6xHis immobilized on Cobalt beads or the same volumes of beads with MBP-6xHis. Beads were pelleted at 16,000 g, and proteins in supernatant were separated by SDS-PAGE, stained with Coomassie, and scanned to measure and calculate the fractions of proteins bound to the beads.

### Co-IP and Western blotting

Co-IP and Western blotting were performed as described except where noted (Laporte et al., 2011; Lee and Wu, 2012; Singh et al., 2024; Ye et al., 2012). Briefly, proteins tagged with mEGFP, mECitrine, GFP, or 13Myc were expressed under the native promotors. Lyophilized cells were ground into a homogeneous fine powder using pestles and mortars. IP buffer (50 mM 4-(2-hydroxyethyl)-1-piperazineethanesulfonic acid [HEPES], pH 7.5, 150 mM NaCl, 1 mM EDTA, 0.1% NP-40, 50 mM NaF, 20 mM glycerophosphate, and 0.1 mM Na_3_VO_4_, 1 mM PMSF, and protease inhibitor [Roche] 1 tablet/30 ml buffer) was added at the ratio of 10 µl: 1 mg lyophilized cell powder. 60 µl Dynabeads protein G beads (Invitrogen) were incubated with 5 µg polyclonal GFP antibody (Novus Bio) for 1 h at room temperature. After three washes with PBS and one wash with 1 ml IP buffer, the beads were incubated with cell lysate for 2 h at 4°C. After 5 washes at 4°C with 1 ml IP buffer each time, the beads were boiled with 80 µl sample buffer. The protein samples were separated with SDS-PAGE gel and detected with monoclonal anti-GFP antibody (1:1,000 dilution; 11814460001; Roche, Mannheim, Germany), monoclonal anti-Myc antibody (1:500 dilution, 9E10, Santa Cruz Biotechnology, Dallas, TX). Secondary anti-mouse immunoglobulin G (1:5,000 dilution; A4416, Sigma-Aldrich) was detected using SuperSignal Maximum Sensitivity Substrate (Thermo Fisher Scientific) on iBright CL1500 imager (Thermo Fisher Scientific) or other imagers.

### Electron microscopy

We grew *ync13Δ* and *ync13^+^* cells exponentially at 25°C in YE5S + 1 M sorbitol for ∼48 h and collected cells by spinning at 2,200 rpm for 3 min. Cells were then washed 2x with equal volume of YE5S medium without sorbitol and diluted to appropriate density and grown 3.5 h at 25°C. OD_600_ of cells was < 0.5 before collecting for high pressure freezing. Sample preparations and electron microscopy were performed as described previously (Kukulski et al., 2012b; Muriel et al., 2021). Briefly, concentrated cell slurry (∼2 µl) was transferred onto specimen carriers (Wohlwend type A, 3 mm wide, 0.1 mm deep) and covered with a flat lid (Wohlwend type B).

The carrier sandwich was immediately processed by high-pressure freezing on a Wohlwend HPF Compact 02. The frozen samples were stored in liquid nitrogen. We opened the carrier sandwich in liquid nitrogen before freeze substitution, which used 1% uranyl acetate in acetone and embedded in Lowicryl HM20 on the Leica AFS 2 robot. Thin sections of 60 nm were cut with a diamond knife using a Leica Ultracut UC7 ultramicrotome and loaded onto carbon-coated 200-mesh copper grids (AGS160; Agar Scientific). The grids were post-stained with 2% uranyl acetate, and then Reynolds lead citrate for 10 min. We observed and imaged the grids using a FEI Tecnai 12 at 120 kV with a bottom mount FEI Eagle camera (4k x 4k).

### Online supplemental material

Fig. S1 shows representative controls for the Tom20-GBP mistargeting experiments. Fig. S2 shows positive and negative physical interactions revealed by mistargeting to mitochondria using Tom20-GBP. Fig. S3 shows physical interactions between full length Ync13 and Rga7 or Rng10 truncations revealed by ectopic mistargeting to mitochondria by Tom20-GBP. Fig. S4 shows localizations of Rga7, Rng10, and Bgs4 in temperature-sensitive mutants. Fig. S5 shows Trs120 accumulates at division site in *ync13Δ*, *rga7Δ*, *rng10Δ*, and *rga7Δ rng10Δ* cells. Videos 1-3 show movements of Trs120-3GFP vesicles to the division site. Online supplemental materials are available at

## Acknowledgements

We thank Juan Carlos, Juan Ribas, and Yajun Liu for yeast strains; Olivia Muriel, Jean Daraspe, and the University of Lausanne Electron Microscopy Facility for help with electron microscopy; Anita Hopper, Jim Hopper, Steve Osmani, and Damien Wilburn for equipment; and current and former members of the Wu and Martin groups for helpful discussions.

Sha Zhang was supported by the Pelotonia Postdoctoral Fellowship Program. J.-Q Wu was supported by fellowships from the Swiss National Science Foundation (grant ISZEZ0_200027) and Herbette Foundation at UNIL to S.G.M. during his faculty professional leave. The work was supported by the National Institute of General Medical Sciences of the National Institutes of Health (grants R01 GM118746 and GM118746-06S1 to J.-Q.W.) and by Swiss National Science Foundation (grant 310030_191990 to S.G.M.). The content is solely the responsibility of the authors and does not necessarily represent the official views of Pelotonia Fellowship Program or The Ohio State University, the National Institutes of Health, or other funding agencies.

## Author contributions

Conceptualization, S.Z., D.S., and J.-Q.W.; Methodology, S.Z., D.S., Y.-H.Z., A.M.-C., and J.-Q.W.; Investigation, S.Z., D.S., Y.-H.Z., K.J.Z., A.M.-C., and J.-Q.W.; Formal Analysis, S.Z., D.S., K.J.Z., J.-Q.W.; Validation, S.Z. D.S.; Visualization, S.Z., D.S., and J.-Q.W.; Writing, S.Z., D.S., S.G.M., and J.-Q.W.; Funding Acquisition, S.Z., S.G.M., and J.-Q.W.; Resources, S.G.M. and J.-Q.W.; Supervision, J.-Q.W.

## Supporting Information

**Table S1.**
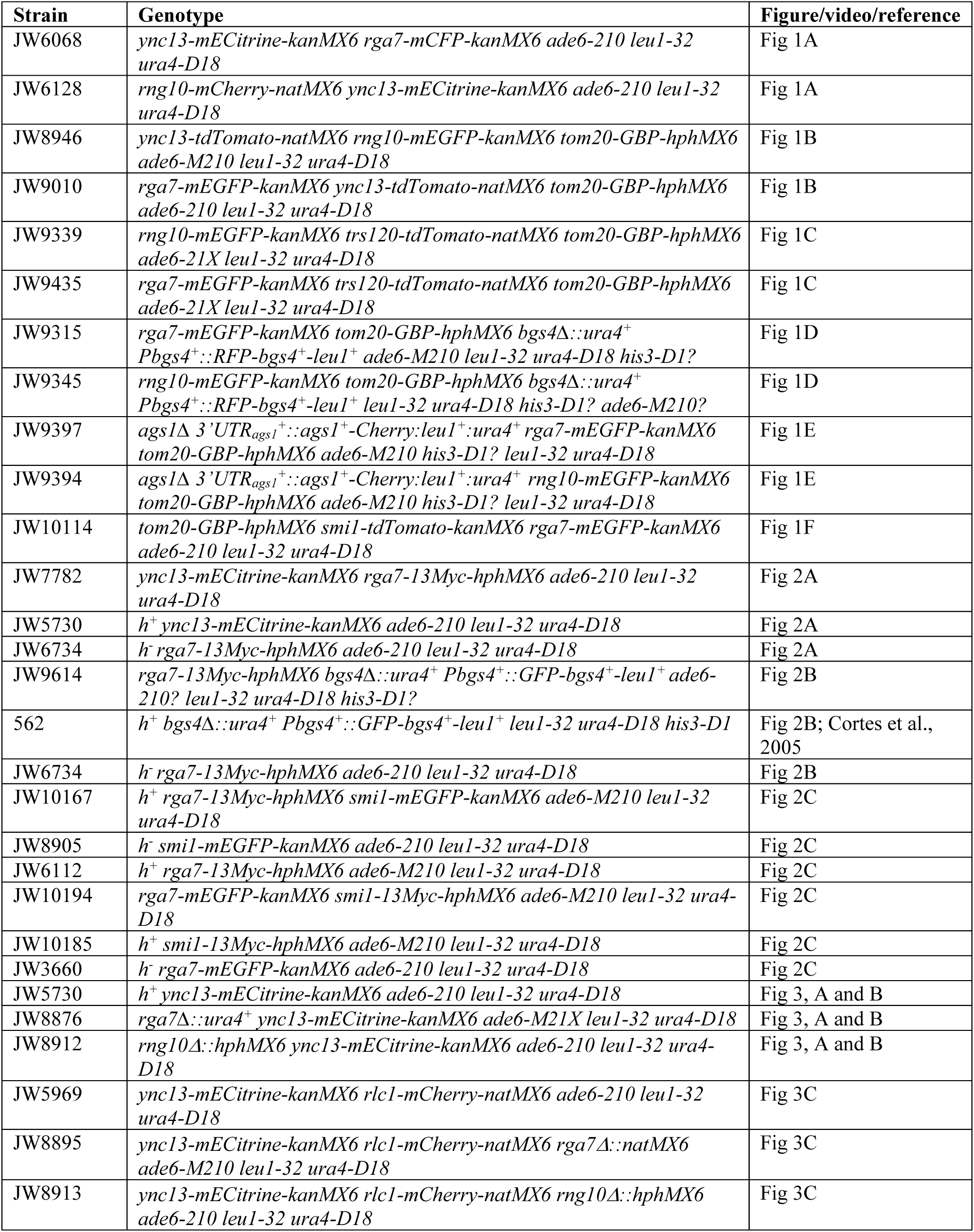
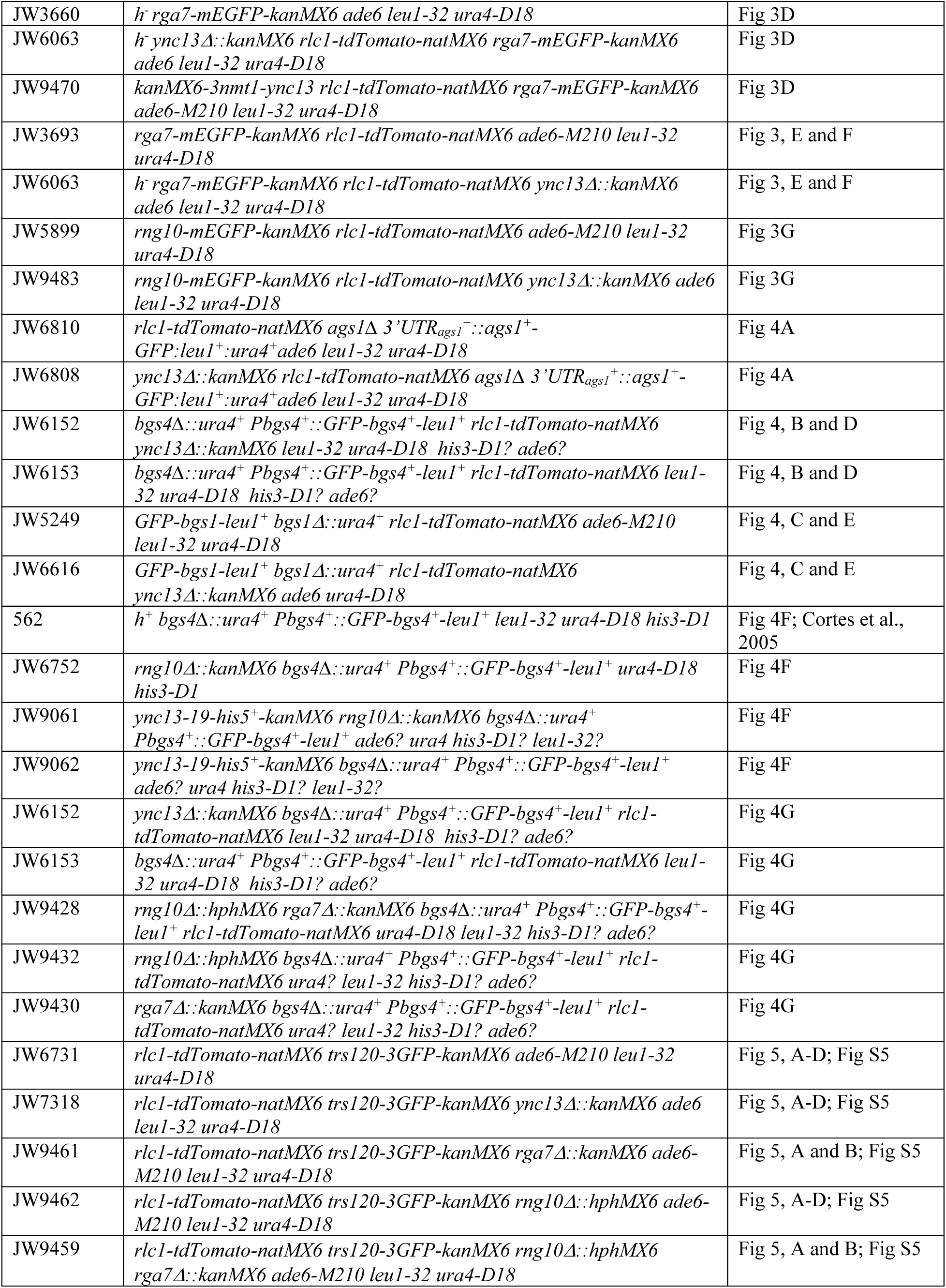
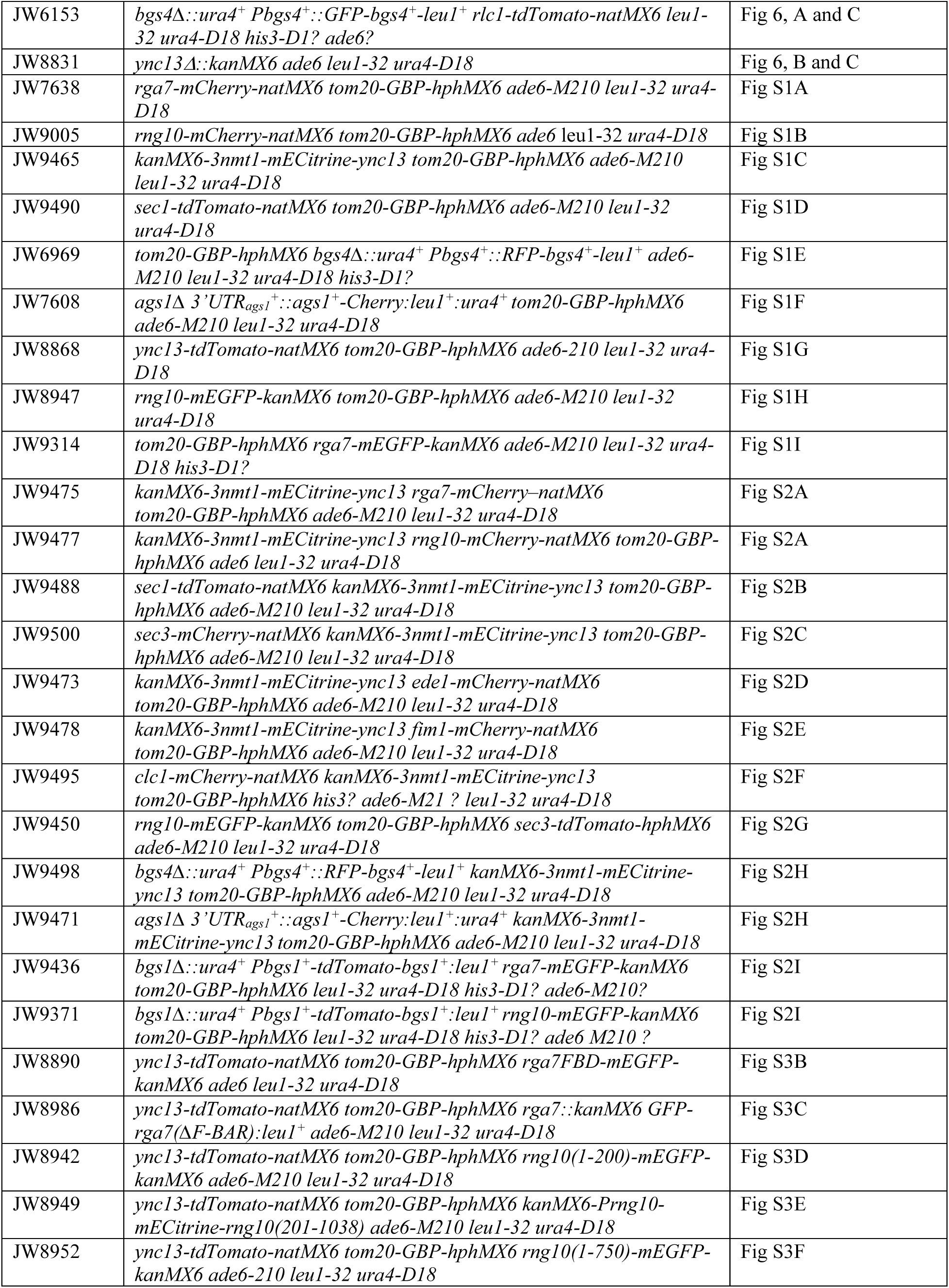
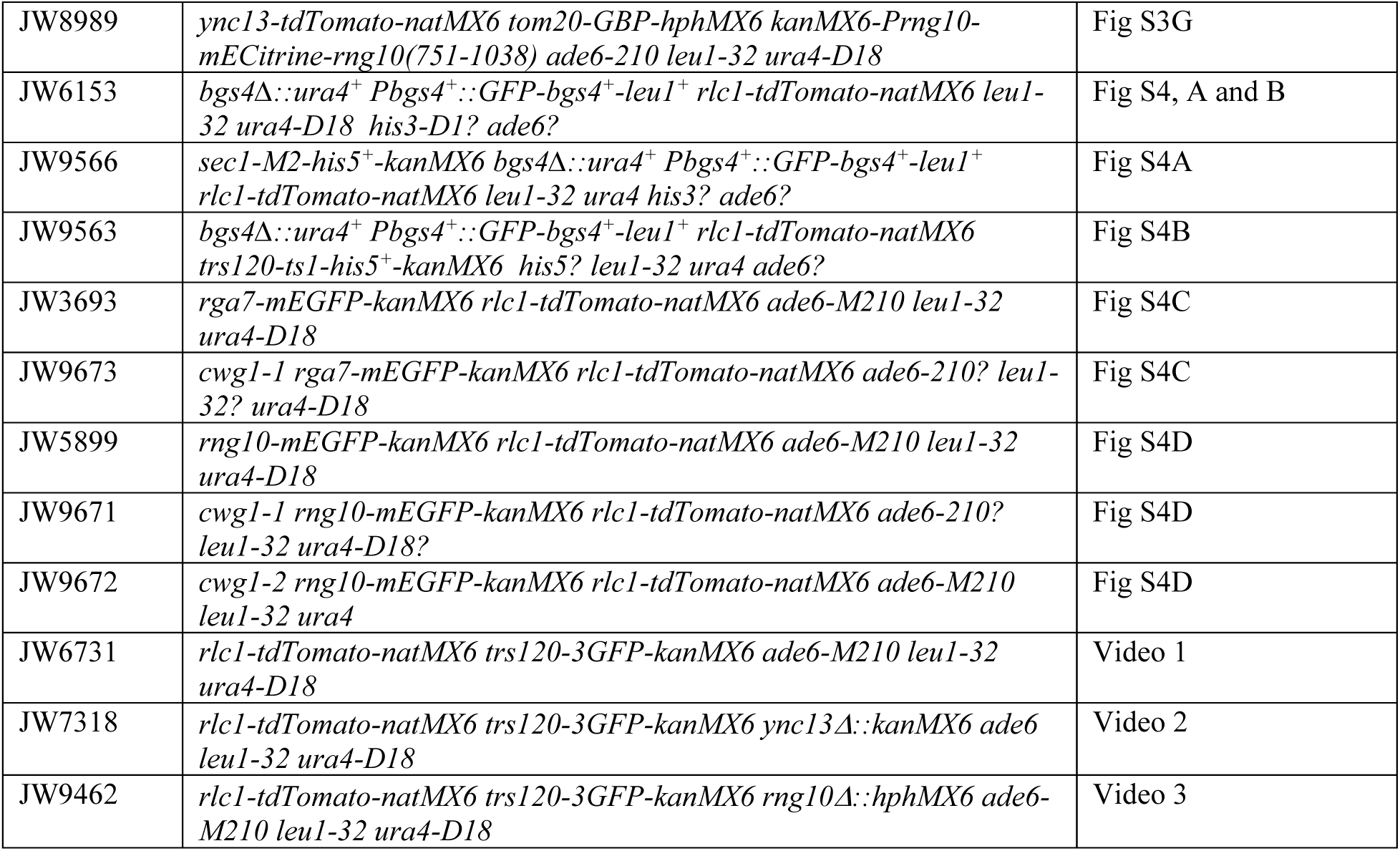
*S. pombe* strains used in this study.

**Figure S1.**
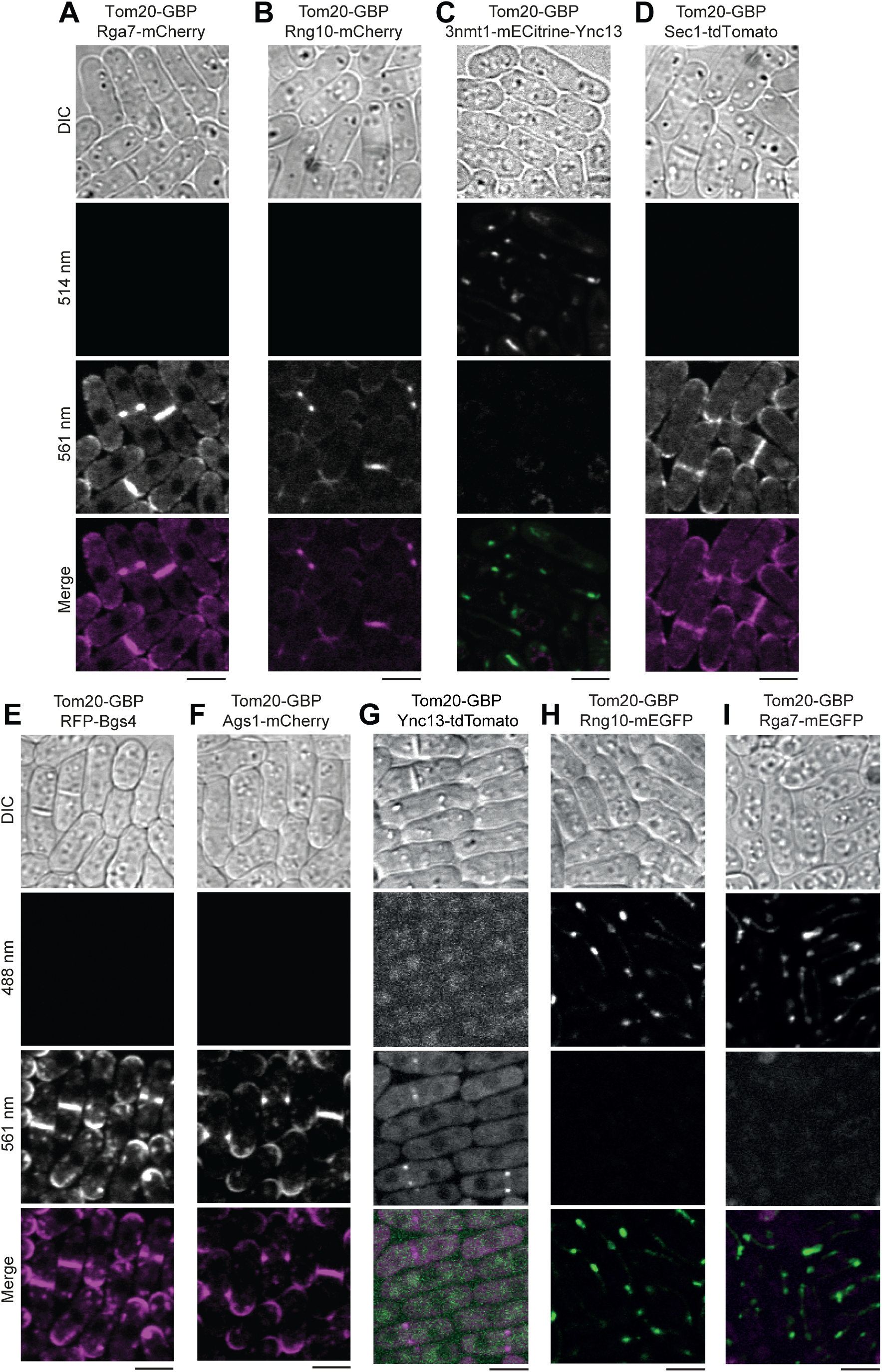
Representative controls for the Tom20-GBP mistargeting experiments. (A-I) Micrographs of DIC, 514/488/561 nm channels, and merged channels showing cells expressing Tom20-GBP and another indicated protein. Tom20-GBP does not bind to mCherry, RFP, or tdTomato so the tagged proteins cannot be recruited to mitochondria without proteins tagged with mEGFP or mECitrine. No signal bleed through between red (561 nm)/yellow (514 nm) or red (561 nm)/green (488 nm) channels. The brightness and contrast were adjusted the same as the experimental groups with three tagged proteins shown in Figs. 1 or S2 so some panels appear almost totally black. Bars, 5 μm.

**Figure S2.**
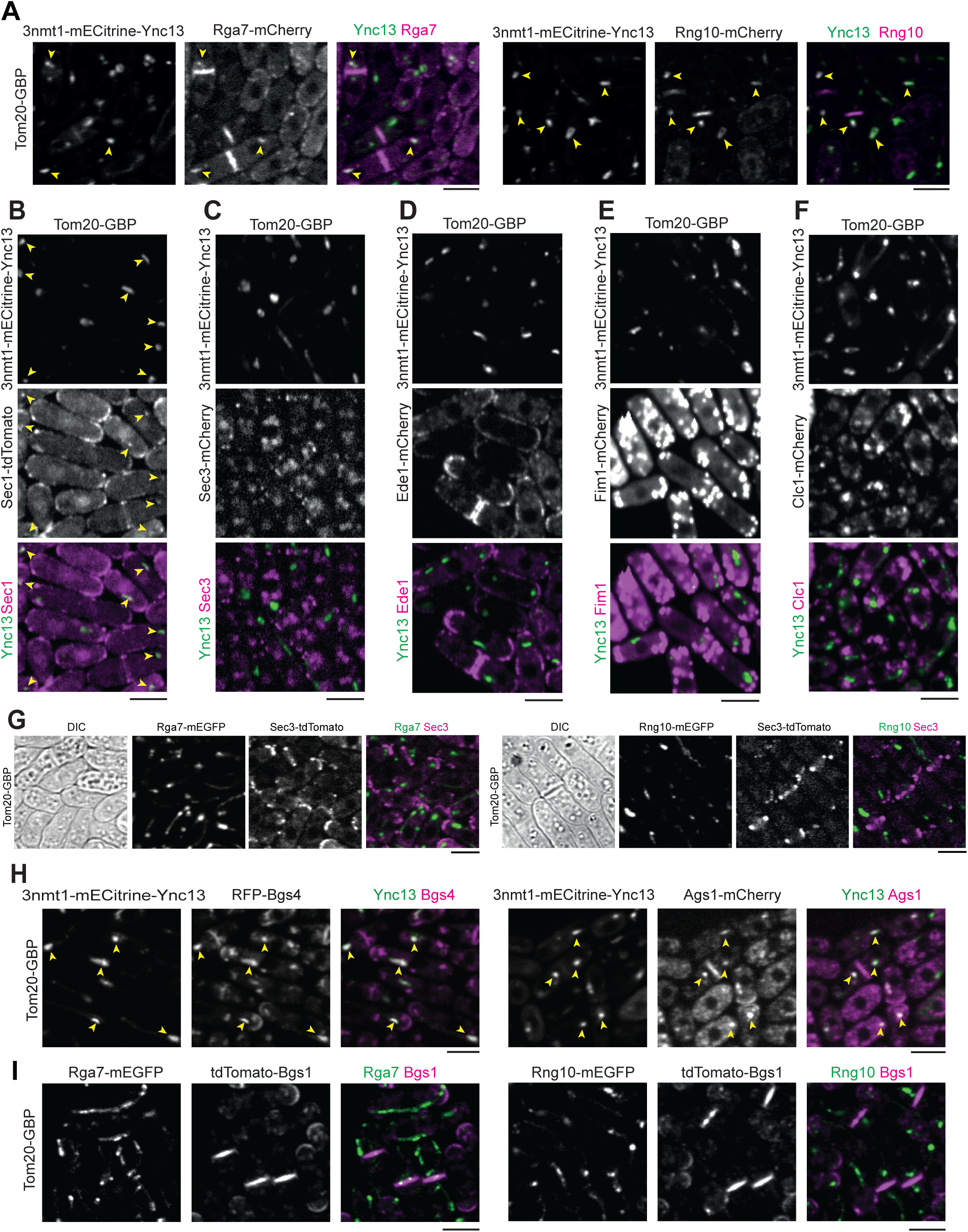
Positive and negative physical interactions revealed by mistargeting to mitochondria using Tom20-GBP. Mislocalized Ync13 ectopically targets (examples marked with arrowheads) Rga7 and Rng10 **(A)**, Sec1 **(B)**, Bgs4 and Ags1 **(H)** to mitochondria, but cannot interact with the exocyst subunit Sec3 **(C)** or proteins Ede1 **(D)**, fimbrin Fim1 **(E),** or clathrin light chain Clc1 **(F)** in the endocytic pathway. Ync13 was overexpressed using the *3nmt1* promoter by growing exponentially in YE5S liquid medium for ∼48 h before imaging. Rga7 and Rng10 cannot mistarget Sec3 **(G)** or Bgs1 **(I)** to mitochondria. Bars, 5 μm.

**Figure S3.**
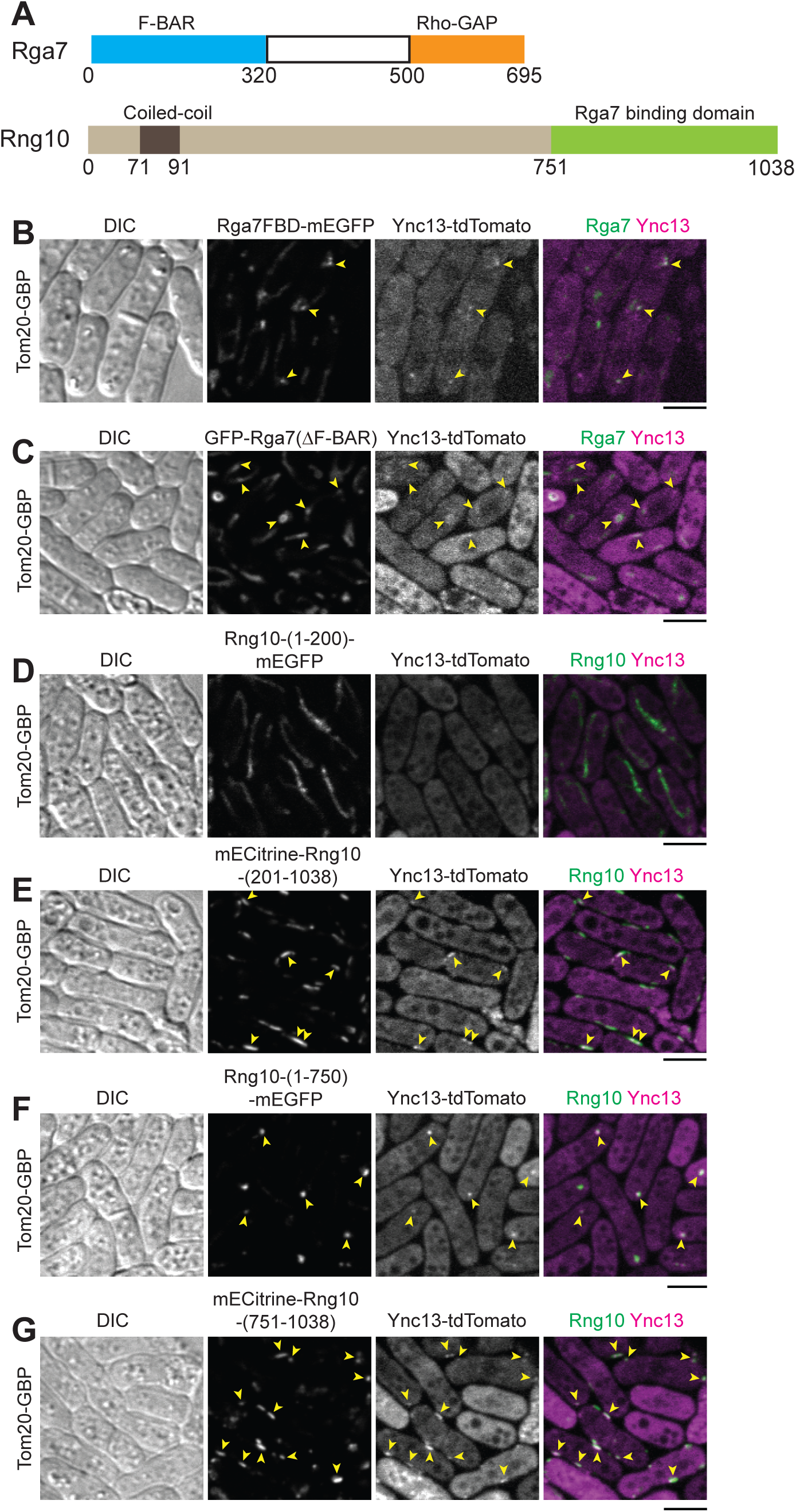
Physical interactions between full length Ync13 and Rga7 or Rng10 truncations revealed by ectopic mistargeting to mitochondria by Tom20-GBP. **(A)** Domain schematics of Rga7 and Rng10 (Liu et al., 2016; Liu et al., 2019). **(B-G)** Arrowheads mark examples of colocalization at mitochondria. Except Rng10-(1-200) in **(D)**, all other Rga7 and Rng10 truncations **(B, C, and E-G)** can mistargeting Ync13-tdToamto to mitochondria. Rga7FBD = Rga7(1-320) (Arasada and Pollard, 2015); Rga7(ΔF-BAR), Rga7 without the F-BAR domain. Bars, 5 μm.

**Figure S4.**
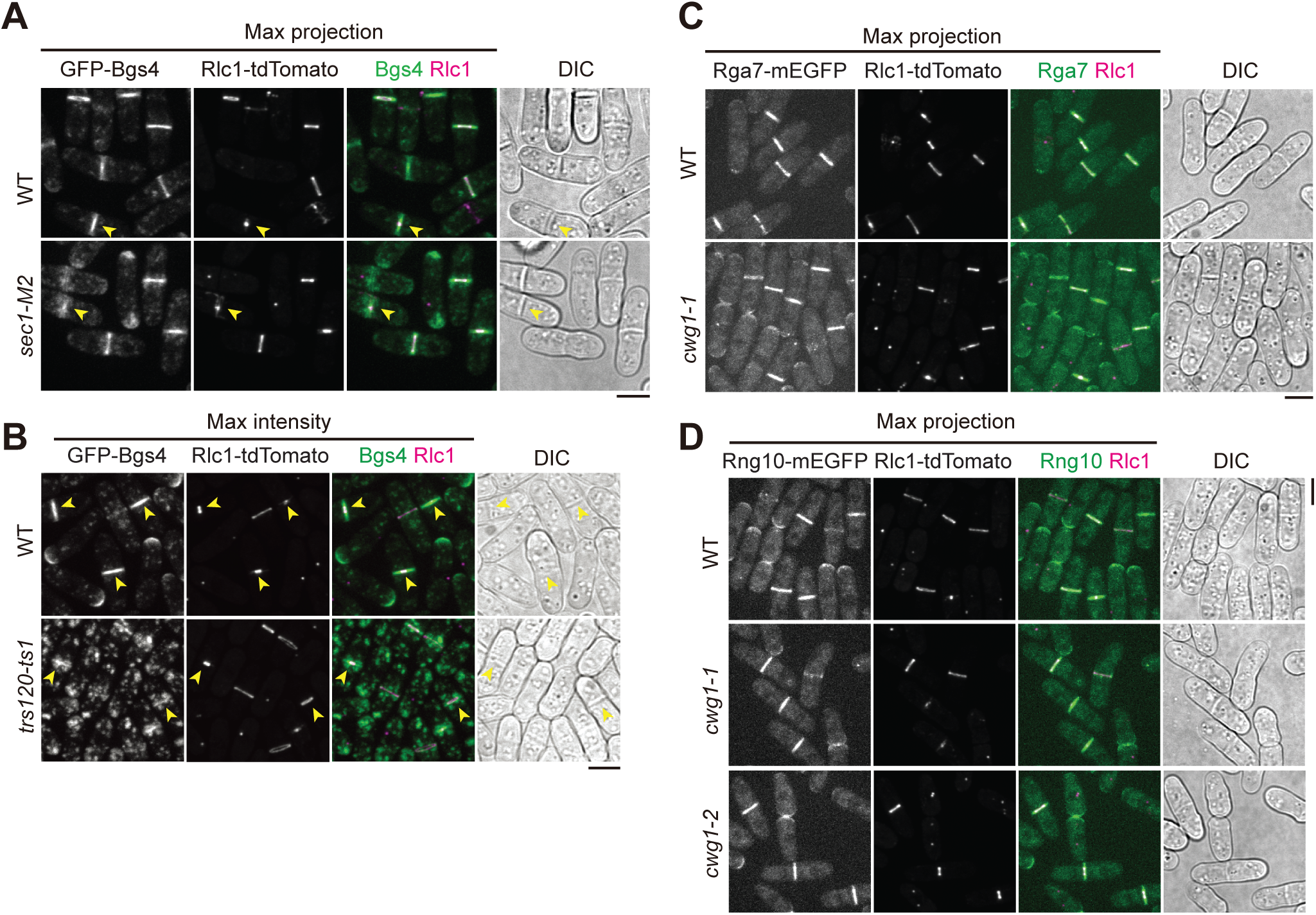
Localizations of Bgs4, Rga7, and Rng10 in temperature-sensitive mutants. Cells grown exponentially at 25°C were shifted to 36°C for 4 h (A, C, D) or 2 h (B) before imaging. Rlc1-tdTomato as the ring marker. Bgs4 localization in *sec1-M2* **(A)** or *trs120-ts1* **(B)** mutant cells. Rga7 **(C)** and Rng10 **(D)** localization in *bgs4* mutants *cwg1-1* and *cwg1-2*. Bars, 5 μm.

**Figure S5.**
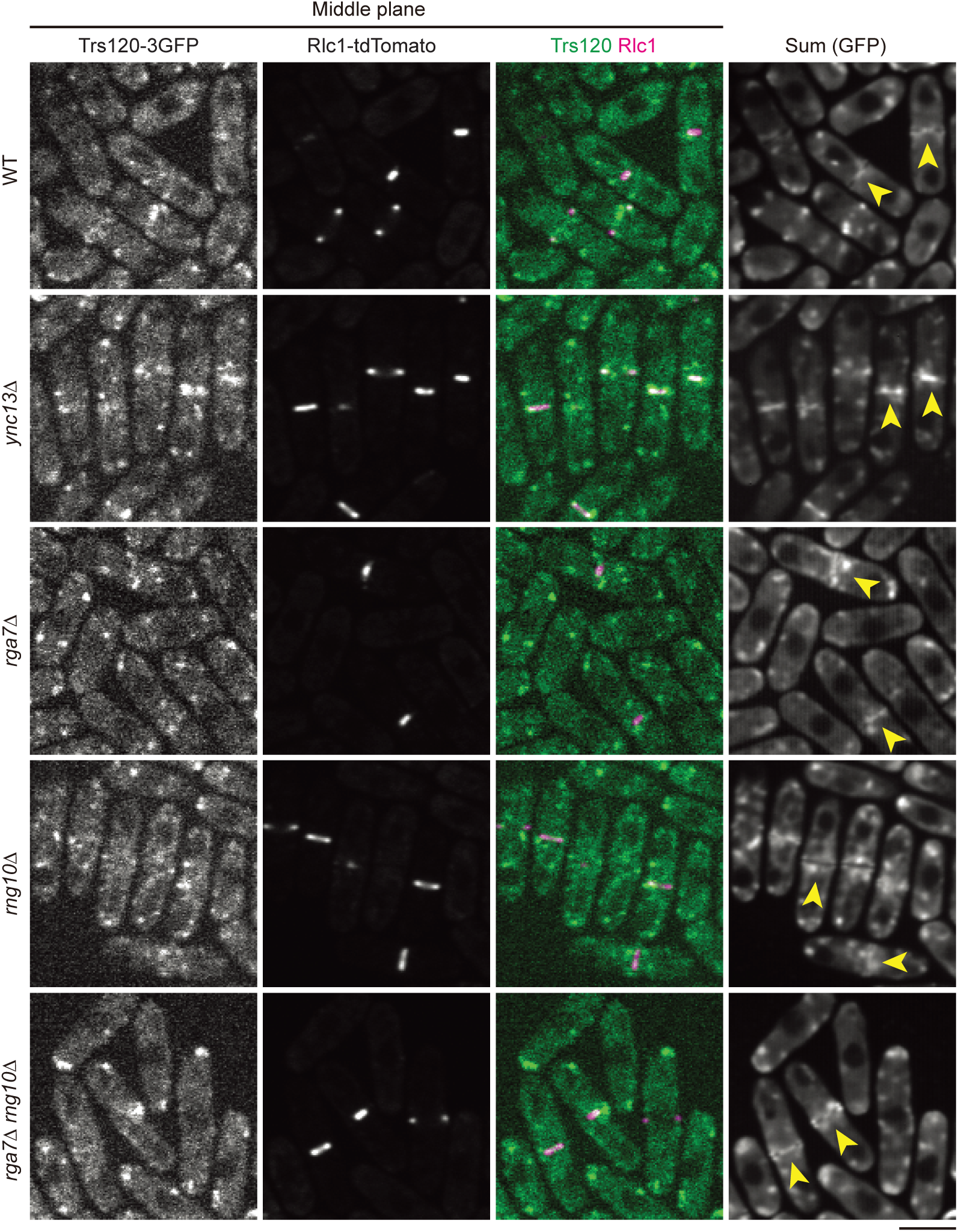
Trs120 accumulates at division site in WT, *ync13Δ*, *rga7Δ*, *rng10Δ*, and *rga7Δ rng10Δ* cells. Rlc1-tdTomato marks the position and diameter of the contractile ring. Trs120-3GFP accumulates inside the area with the ring in *ync13Δ* cells, but outside the ring area in *rga7Δ*, *rng10Δ*, and *rga7Δ rng10Δ* cells at division site. The middle focal planes along with the sum intensity projection of the GFP channel from a 2-min continuous movie (exposure time 200 ms) without delay are shown. Arrowheads mark Trs120-3GFP in cells with constricting ring. Cells were grown exponentially in EMM5S liquid media for ∼48 h before imaging. Bars, 5 μm.

## Movie legends

**Videos 1-3. Movement of Trs120-3GFP vesicles to the division site.** The movies track the movement of Trs120-3GFP vesicles from the cytoplasm to the division site in WT (Video 1), *rng10*Δ (Video 2), and *ync13*Δ (Video 3) cells using colored lines. The videos correspond to Fig. 5 and Fig. S5. Strains were grown exponentially in EMMS liquid media at 25°C for ∼48 h before imaging in continuous movies without delay. Time, min: sec.

